# Chemical Dice Integrator (CDI): A Scalable Framework for Multimodal Molecular Representation Learning

**DOI:** 10.1101/2025.11.11.687860

**Authors:** Suvendu Kumar, Saveena Solanki, Mudit Gupta, Sanjay Kumar Mohanty, Shiva Satija, Sonam Chauhan, Subhadeep Duari, Arushi Sharma, Vishakha Gautam, Sakshi Arora, Raidhani Shome, Sourav Sinha, Abhinav Kumar Sharma, Aayushi Mittal, Debarka Sengupta, Natarajan Arul Murugan, Gaurav Ahuja

## Abstract

The machine learning landscape for molecular property prediction is fragmented, with numerous Featurizers each capturing a narrow, specialized view of chemical structure. This heterogeneity forces a suboptimal choice of representation a priori, limiting model generalizability. We introduce the Chemical Dice Integrator (CDI), a hierarchical framework that unifies six orthogonal molecular representations, physicochemical (Mordred), topological (GROVER), visual (ImageMol), biological (Signaturizer), quantum-mechanical (MOPAC), and linguistic (ChemBERTa), into a single, coherent embedding. The framework consists of CDI-Basic, a two-tiered autoencoder that fuses these modalities, and CDI-Generalised, a Mamba State-Space Model (SSM) that learns a direct, efficient map from SMILES strings to the unified embedding space. Extensive benchmarking across 23 classification (171 tasks) and 10 regression datasets demonstrates that CDI embeddings consistently achieve superior predictive performance compared to individual Featurizers and standard feature aggregation methods. The CDI-Generalised model achieves this performance with exceptional computational efficiency, outperforming deep learning Featurizers in terms of speed and resource overhead. Furthermore, we demonstrate that the CDI embedding is chemically intuitive, allowing for the sensitive distinction of nuanced structural variants, such as chiral enantiomers and kekulized SMILES forms. By bridging multimodal chemical intelligence with scalable, sequence-based inference, CDI offers a strong foundation for molecular machine learning.

## INTRODUCTION

The digitization of molecular structure is a foundational step in applying machine learning (ML) to chemical and pharmaceutical research, with the choice of representation having a critical influence on model performance ^1^. The field has witnessed an evolution from expert-curated physicochemical descriptors ^2^ and fingerprints to quantum mechanical simulations capturing electronic properties ^3^. The deep learning revolution further expanded this toolkit, introducing powerful, data-driven representations that reframe the molecule as a graph ^4^, a string ^5^, or a 2D image ^6^. Each of these featurization strategies embodies a different inductive bias, leading to significant gains on specific benchmarks where their particular view of chemical space is advantageous. However, this proliferation of specialized Featurizers has created a critical paradox: an abundance of choice that now hinders generalizability ^7,8^. No single representation, be it the topological awareness of graph networks (GROVER ^4^), the syntactic understanding of language models (ChemBERTa ^5^), or the bioactivity profiles from specialized predictors (Signaturizer ^9^), consistently dominates across the vast and diverse spectrum of prediction tasks in drug discovery ^10^. This performance fragmentation forces researchers into a costly and inefficient cycle of task-specific Featurizer selection, which is often based on heuristic experience rather than principled methodology. Consequently, this undermines the development of robust, universal predictive models that can generalize across different endpoints, from predicting pharmacokinetics (ADMET) and toxicity to quantifying quantum interactions.

Current attempts to integrate these disparate modalities often rely on simplistic methods such as feature concatenation or the application of generic dimensionality reduction techniques like Principal Component Analysis (PCA). These approaches are demonstrably insufficient because they operate under a flawed assumption: that the complex, non-linear relationships between orthogonal chemical modalities exist on a linear or easily projectable manifold. For instance, they fail to capture the profound interconnections between a molecule’s quantum mechanical landscape (e.g., as determined by MOPAC calculations ^11^) and its macroscopic bioactivity profile (e.g., as determined by Signaturizer ^9^). This limitation highlights a pressing and unmet need for a new paradigm in molecular representation learning, one that transcends mere combination to achieve true integration. The field requires a learned, unified representation that can holistically capture the multifaceted nature of chemical compounds.

Here, we introduce the Chemical Dice Integrator (CDI), a hierarchical deep learning framework designed to overcome this fundamental fragmentation. CDI represents a paradigm shift from ensemble-based or concatenative approaches by constructing a foundational, holistic molecular embedding through intelligent, hierarchical fusion. The framework is a game-changer for two core reasons. First, its CDI-Basic engine is a novel multimodal fusion architecture that employs a two-tiered autoencoder to seamlessly integrate six orthogonal representations, spanning physicochemical (Mordred ^2^), topological (GROVER ^4^), visual (ImageMol ^6^), biological (Signaturizer ^9^), quantum (MOPAC ^11^), and linguistic (ChemBERTa ^5^) domains into a single, coherent embedding space. This process does not merely compress data; it actively learns the latent semantic structure that interconnects these disparate views of a molecule, capturing the synergistic information that exists between them. Second, and critically for scalability, we developed CDI-Generalised. This model incorporates a modern Mamba State-Space Model (SSM) ^12^ to learn a direct, highly efficient mapping from a raw SMILES string to the unified CDI embedding space. This innovation bypasses the computational bottleneck of generating all six source features during inference, a key limitation that has previously hampered the practical deployment of multimodal approaches. Through rigorous and extensive benchmarking across a vast suite of 23 classification (171 tasks) and 10 regression datasets, we demonstrate that CDI not only matches but frequently outperforms its constituent Featurizers and other integration baselines. Furthermore, we provide evidence that the resulting CDI embedding is chemically intuitive, capable of distinguishing subtle structural nuances, such as chiral centers and kekulization states, that elude other methods. By solving the fragmentation problem at the representation level and providing a scalable inference pipeline, the CDI framework establishes a new, production-ready foundation for molecular machine learning, with the potential to significantly accelerate the pace of discovery in therapeutics and materials science.

## RESULTS

### A Unified Molecular Embedding Framework via Multimodal Fusion and Sequence-Based Inference

We developed the Chemical Dice Integrator (CDI) to generate a universal molecular representation that is both chemically comprehensive and computationally scalable. The CDI framework is built on a dual-component architecture: a multimodal fusion engine (CDI-Basic) and a sequence-to-embedding model (CDI-Generalised). The CDI-Basic engine integrates six orthogonal molecular representations into a single, 8192-dimensional embedding **(Figure 1a)**. Rather than concatenating features, we employed a two-tiered hierarchical autoencoder architecture designed to learn deep semantic relationships between modalities. In the first tier, six Semantic Commonality Autoencoders (SCAs) are trained using a leave-one-out reconstruction objective, where each autoencoder learns to infer a shared latent subspace from the other five modalities **(Figure 1c, left)**. This process enforces the discovery of shared inter-modality information and yields six latent subspaces that capture semantic overlap among feature domains. In the second tier, the outputs are then fused by a Super-Embedding Autoencoder (SEA) to produce the final CDI-Basic embedding, which produces the unified CDI-Basic embedding while minimizing reconstruction and semantic alignment losses. The full mathematical formulation and implementation details of this architecture, including reconstruction and mean square error loss functions and encoder-decoder mappings, are provided in **Supplementary Figure 1a**. This fusion process demonstrated stable convergence over 600 epochs, confirming stable multimodal integration during training **(Figure 1e)**. The dataset for this pre-training was rigorously curated from the ChEMBL (v35) corpus, with a detailed preparation workflow shown in **(Supplementary Figure 1b)**.

**Figure 1:**
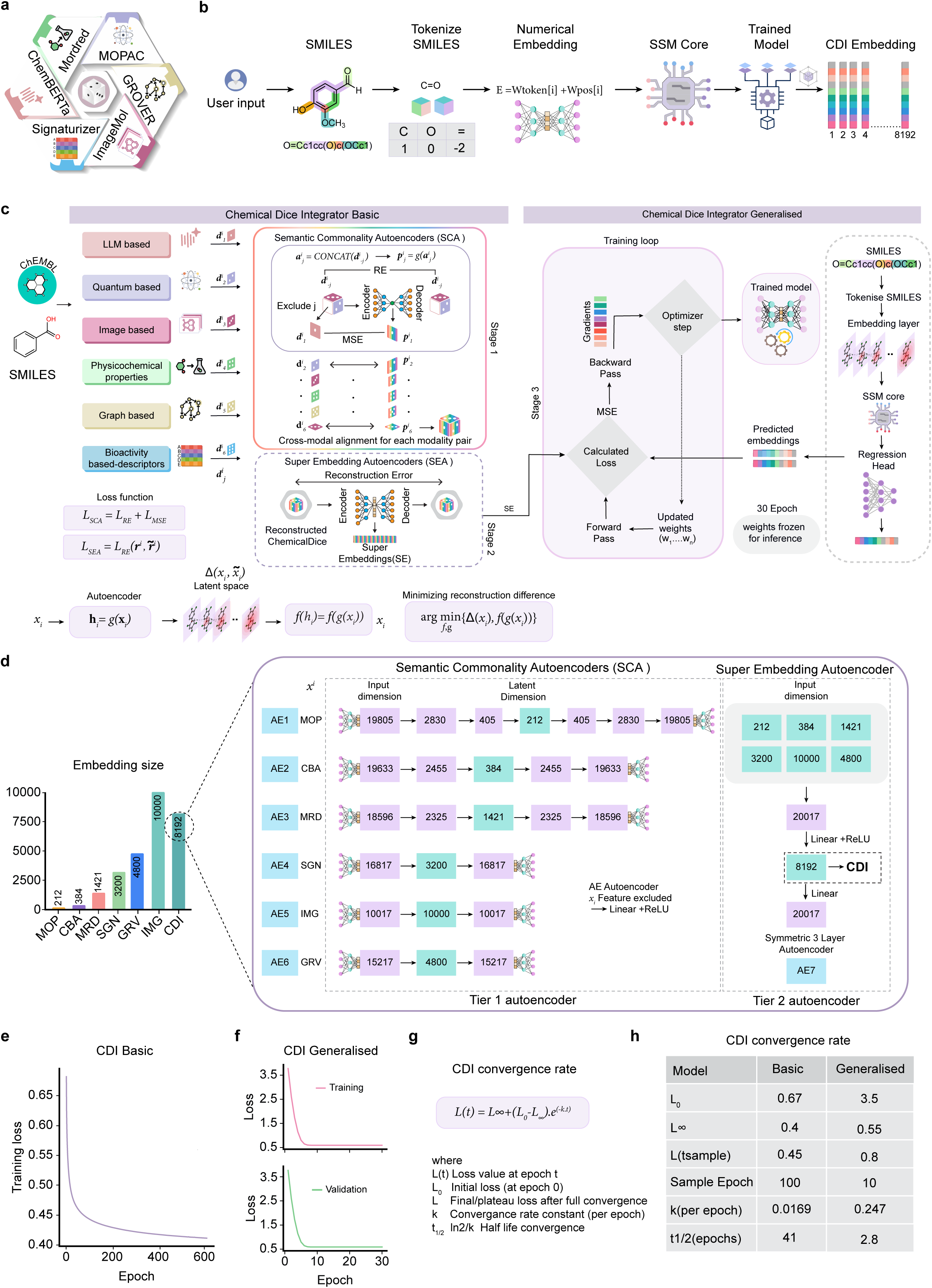
The Chemical Dice Integrator (CDI) Architecture and Initial Validation. **(a)** Schematic overview of the six complementary molecular representations integrated by the CDI framework, encompassing physicochemical (Mordred), graph-based (GROVER), image-based (ImageMol), bioactivity (Signaturizer), quantum-mechanical (MOPAC), and language-based (ChemBERTa) embeddings. **(b)** The inference workflow for CDI-Generalised, depicting the pathway from a user-input SMILES string through tokenization, an input embedding layer, the Mamba State-Space Model (SSM) core, and a final regression head to produce the predicted CDI embedding. **(c)** The end-to-end training pipeline illustrates the construction of CDI-Basic and CDI-Generalised. The ChEMBL dataset generates the six source representations. The CDI-Basic model is built via a two-tiered autoencoder: first, six Semantic Commonality Autoencoders (SCAs) are trained using a leave-one-out reconstruction strategy, and their latent outputs are fused by a Super-Embedding Autoencoder (SEA) to create the unified CDI-Basic embedding. These embeddings then serve as regression targets for training the SMI-SSED (SMILES-based State-Space Encoder-Decoder) model to create CDI-Generalised. **(d)** A bar plot depicting the embedding sizes across modalities, along with their respective layer configurations and the latent dimensional hierarchies of SCAs and SEA that culminate in the final CDI embedding. **(e)** The training loss curve for the CDI-Basic fusion autoencoder is over 600 epochs, demonstrating stable convergence. **(f)** The training (top) and validation (bottom) loss curves for the CDI-Generalised model over 30 epochs, indicating effective learning without overfitting. **(g)** Mathematical formulation of CDI’s convergence kinetics, modeled as an exponential decay function. **(h)** Comparative convergence parameters for CDI and CDI-Generalised models. The Generalised variant exhibits a markedly higher convergence rate constant (k = 0.247 epochL¹) and a shorter half-life (tL/L = 2.8 epochs), demonstrating faster stabilization and more efficient optimization relative to the CDI (Chemical Dice Integrator) model.

To enable scalable inference, we developed CDI-Generalised, which replaces the multimodal preprocessing step with a direct sequence-to-embedding translation mechanism. This model leverages a Mamba State-Space Model (SSM) core as a direct sequence-to-embedding mapper (SMI-SSED), trained to predict the CDI embedding space directly from tokenized SMILES **(Figure 1b, c, right)**. A comparison of embedding dimensionalities shows CDI’s representation is both compact and information-dense relative to its constituent Featurizers **(Figure 1d)**. CDI-Generalised was trained using a composite loss function combining mean-squared and cosine similarity components, ensuring both geometric and angular alignment with the CDI-Basic target embeddings. The model was optimized with a composite loss, resulting in robust training dynamics without overfitting across 30 epochs **(Figure 1f)**. Quantitative convergence metrics **(Figures 1g-h)** show that CDI-Basic and Generalised achieved a 15-fold faster convergence rate (half-life ≈ 2.8 epochs) and required nearly ten-fold fewer parameters (0.108 B vs 1.09 B) than CDI-Basic. Hyperparameters and training settings for both CDI variants are summarized in **(Supplementary Figure 1c)**, showing that CDI-Generalised achieves high fidelity with substantially fewer parameters and epochs. This approach effectively decouples the rich, multimodal representation from the computational cost of generating all six source Featurizers during inference. This architecture allows CDI to preserve the semantic depth of multimodal integration while achieving the scalability necessary for high-throughput analysis.

### The CDI Embedding Outperforms Traditional Feature-Set Aggregation Methods

To determine whether the unified CDI embedding provides a fundamental advantage over traditional feature aggregation techniques, we designed a rigorous, large-scale benchmarking suite termed the "Aggregator" module. This module evaluates CDI against a comprehensive panel of eight well-established linear and non-linear dimensionality reduction methods, including Principal Component Analysis (PCA), Canonical Correlation Analysis (CCA), Independent Component Analysis (ICA), Kernel PCA (kPCA), Random Kitchen Sinks (RKS), Isomap, Locally Linear Embedding (LLE), and t-distributed Stochastic Neighbor Embedding (t-SNE) **(Figure 2b)**. Aggregation here refers to the process of projecting heterogeneous feature spaces into a common latent manifold using kernel or projection-based transformations such as PCA or RKS. The evaluation spanned a vast chemical space, comprising 23 classification datasets (totaling 171 distinct tasks) and 10 regression datasets, covering critical endpoints in toxicity (AMES, DILI, Tox21), ADMET (PGP, BBB, CYP450 inhibition), and physicochemical properties (Solubility, Lipophilicity, clearance) **(Figure 2a)**. The results demonstrate the unequivocal superiority of the CDI embedding. When evaluated using a uniform pipeline of nine distinct machine learning models, CDI achieved the highest mean scores across critical metrics, including AUC-ROC and Balanced Accuracy, significantly outperforming all other Aggregator sets **(Figure 2c)**. Its performance advantage was robust across model architectures, confirming that CDI captures a universally informative feature manifold rather than one tailored to a single algorithm. Bar plots with standard error of mean confirmed that CDI delivered the top mean scores with the smallest confidence widths, evidencing tight generalization and reproducible behavior **(Figure 2d)**. The superiority of CDI was not an artifact of a single model, but rather a robust trend, as visualized by performance radar charts revealing CDI’s dominance across seven key metrics: Accuracy, Precision, Recall, F1-Score, Cohen’s Kappa, Balanced Accuracy, and AUC-ROC. CDI formed the outermost envelope of the radar chart, indicating that its latent space supports balanced performance across all predictive dimensions **(Figures 2c and h)**. This demonstrates that the CDI embedding provides a more generically useful feature space for a wide range of algorithms. A comprehensive statistical comparison using raincloud plots, which combine violin, box, and jittered point distributions, confirms that CDI shows the most consistent (narrowest) performance distribution for both AUC-ROC and Balanced Accuracy, indicating high reliability and uniform predictive behavior across tasks. Pairwise Mann-Whitney U tests were applied to compare median performance distributions between CDI and all baseline aggregation methods across each evaluation metric (AUC-ROC, Balanced Accuracy, Accuracy, Precision, Recall, F1-Score, and Kappa). To correct for multiple comparisons, p-values were adjusted using the Bonferroni false-discovery rate (FDR) correction, ensuring stringent control of Type I errors. Statistical significance thresholds were set as p < 0.05 (*), p < 0.01 (**), and p < 0.001 (***), as indicated above the violin plots **(Supplementary Figure 2a-b)**. Bar plots of mean performance with 95% confidence intervals further underscore CDI’s significant and consistent advantage, with the highest mean scores and narrowest confidence intervals, confirming its better and consistent generalization performance across diverse datasets **(Supplementary Figure 2c; Supplementary Table 4)**. The scale of CDI’s dominance is visually striking when examining the performance differentials (Δ) between CDI and the next-best method ("Winner") across hundreds of model-task combinations. Contour density jitter plots reveal a dense cloud of data points tightly clustered near zero on both axes for Δ Accuracy versus Δ Precision **(Figure 2e)** and Δ Recall versus Δ F1-Score **(Figure 2f)**, indicating that CDI was most frequently the top performer. Similarly, the cloud for Δ Balanced Accuracy versus Δ AUC-ROC is sharply concentrated at the origin **(Figure 2g)**, underscoring that when CDI is not the absolute winner, its performance is negligibly close. This comprehensive superiority is further synthesized in a radial plot, where CDI forms the outermost envelope across all seven performance metrics, including Precision,Recall, where CDI formed the outermost and most balanced performance envelope across all metrics, including Kappa, underscoring its robustness and uniform predictive utility **(Figure 2h)**. Pairwise statistical evaluation using one-way ANOVA confirmed that the CDI embedding formed a statistically distinct high-performing population across both AUC-ROC and Balanced Accuracy metrics **(Figure 2i).** Post-hoc Bonferroni-FDR corrected comparisons revealed significant mean differences between CDI and all other aggregation methods, with consistently positive effect sizes. The heatmap distribution illustrates that CDI maintained the largest positive mean shift relative to other feature sets, while inter-baseline comparisons showed comparatively minor or nonsignificant differences. These findings establish that the performance advantage of CDI is statistically robust and intrinsic to its representational quality rather than dependent on model-specific bias. To further validate these findings, a one-way ANOVA followed by Tukey’s HSD post-hoc test (Bonferroni-corrected) was conducted across all feature representations **(Supplementary Fig. 2d)**. The resulting heatmaps summarize mean differences for Accuracy, Balanced Accuracy, Recall, F1-score, Precision, and Kappa, confirming that CDI consistently achieved the highest mean performance and statistically distinct group separation across all evaluated dimensions. To dissect the source of this performance gap, we conducted a multi-factorial ANOVA. The analysis identified the feature set as the dominant factor influencing model output, with an effect size that dwarfed that of the choice of machine learning model or their interaction **(Figure 2j, k)**. The F-statistic for the feature set factor was orders of magnitude greater than for other factors **(Figure 2j)**, and one-way ANOVA specifically on AUC-ROC and Balanced Accuracy again highlighted CDI as a statistically distinct population **(Figure 2i)**. This conclusively demonstrates that the performance gain is inherent to the CDI embedding itself, which captures a more predictive and generalizable chemical feature space than any traditional aggregation method can construct. The statistical conclusions are further reinforced by pairwise comparison heatmaps based on Tukey’s Honest Significant Difference (HSD) post-hoc test, which show CDI consistently has significant positive mean differences over all other feature sets across all metrics **(Supplementary Figure 2d)**. Extended benchmarking analyses **(Supplementary Figures 3-4)** evaluated the statistical robustness of CDI across nine machine-learning models and eight feature aggregation methods. Each model-Aggregator combination was assessed using boxplots for Precision **(Supplementary Fig. 3a)**, Recall **(Supplementary Fig. 3b)**, AUC-ROC **(Supplementary Fig. 3c)**, Cohen’s Kappa **(Supplementary Fig. 4a)**, Balanced Accuracy **(Supplementary Fig 4b)**, and Accuracy **(Supplementary Fig. 4c)**. Pairwise Mann-Whitney U tests with Bonferroni FDR correction were performed to compare the distributions of each feature set across models, with significance. These results consistently demonstrated CDI’s statistically significant advantage in all metrics relative to competing methods, with minimal inter-model variance.

**Figure 2:**
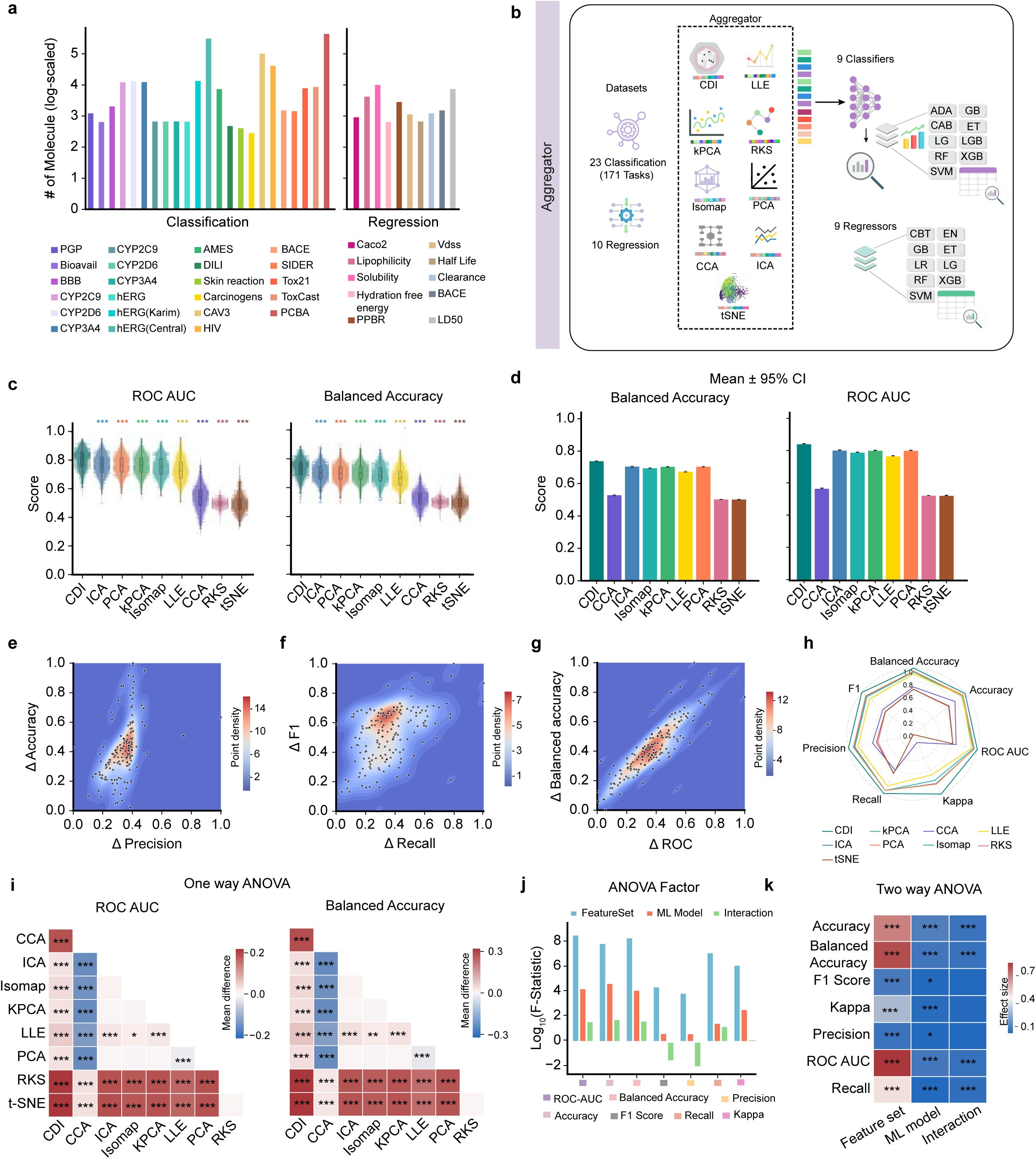
Large-Scale Benchmarking Demonstrates the Superior Predictive Performance of the CDI Embedding. **(a)** Bar chart (log-scaled y-axis) illustrating the number of molecules in each of the 23 classification and 10 regression datasets constituting the Aggregator and Featurizer benchmark suite, covering endpoints in toxicity (AMES, DILI, Toxcast, Tox21), ADMET (PGP, BBB, CYP450 inhibition), and physicochemical properties (Solubility, Lipophilicity, Hydration free energy, Half-life, Clearance, LD50). **(b)** Schematic diagram depicting the Aggregator benchmarking strategy, where all datasets are encoded using CDI and eight other feature aggregation/dimensionality reduction methods (PCA, CCA, ICA, kPCA, RKS, Isomap, LLE, t-SNE), and subsequently evaluated through a uniform pipeline of nine classifiers and nine regressors, thereby ensuring model-independent comparison. **(c)** Violin distributions representing the overall performance (AUC-ROC and Balanced Accuracy) of CDI and other aggregation methods across all tasks, highlighting CDI’s consistently higher mean and narrower variance. mean AUC-ROC and Balanced Accuracy for all feature sets across all tasks,asterisks denote statistical significance (Mann-Whitney U test) for CDI versus the other method. **(d)** Mean performance with 95% confidence intervals for AUC-ROC and Balanced Accuracy across all datasets, underscoring the significant and consistent advantage of the CDI embedding. **(e)** Contour jitter plots illustrating the performance differentials (ΔAccuracy vs ΔPrecision) between CDI and the top-performing model across hundreds of model–dataset combinations, showing dense clustering near the origin and confirming CDI’s frequent dominance. **(f)** Contour density map of ΔF1 vs ΔRecall visualizing CDI’s balanced improvement in sensitivity and predictive precision relative to competing aggregation techniques. **(g)** Contour plot of ΔBalanced Accuracy vs ΔAUC-ROC displaying CDI’s near-zero deviation from or consistent outperformance of the best baseline method, underscoring its generalizability across classification tasks. **(h)** Radar plot synthesizing the performance profiles of all feature sets across seven key metrics, with CDI forming the outermost and thus superior profile. **(i)** One-way ANOVA comparison across all feature aggregation methods for AUC-ROC and Balanced Accuracy demonstrating that CDI forms a statistically distinct high-performing group (***p < 0.001) relative to all other techniques. A one-way ANOVA was used to evaluate the effect of feature representation on performance metrics, followed by Tukey’s HSD post-hoc test to identify statistically significant pairwise differences between feature sets, with multiple-testing correction. **(j)** Bar plot of the log_10_ (F-statistic) from a two-way ANOVA, identifying the feature set as the dominant factor dictating model performance. **(k)** Effect size (Partial Eta Squared) from the two-way ANOVA for the factors of Feature Set, ML Model, and their Interaction across all performance metrics, reaffirming the paramount importance of the feature set. A two-way ANOVA quantified variance contributions from feature set, machine learning model, and their interaction, confirming that the feature representation factor predominantly influences performance across metrics. Statistical significance was set at pL<L0.05, with significance levels denoted as *L<L0.05, **L<L0.01, and ***L<L0.001.

Extended benchmarking across regression tasks revealed CDI’s versatility. Across model-wise comparisons of 9 metrics shown as heatmaps of classification datasets **(Supplementary Fig. 5a)**, CDI consistently ranked among the top-performing feature representations across all nine classifiers, including both ensemble-based and kernel-based algorithms. Boosting frameworks such as LightGBM, XGBoost, and Gradient Boosting achieved the most stable and highest-ranked performance with CDI embeddings, indicating strong compatibility between CDI’s unified feature space and ensemble learning paradigms. This trend highlights CDI’s robustness and adaptability across diverse machine-learning architectures, further supporting its role as a general-purpose molecular representation. Heatmaps of regression performance across datasets and models show that tree-based and boosting regressors achieve the highest CDI ranks (closest to 1) across R², MAE, RMSE, and MSE metrics **(Supplementary Figure 6a)**. CDI demonstrates the highest median R² across all regressors, confirming superior explanatory power and feature-target alignment **(Supplementary Figure 6b)**. Furthermore, CDI consistently achieves the lowest RMSE **(Supplementary Figure 6c)** and the lowest Mean Absolute Error (MAE) and Mean Squared Error (MSE) values across all regressors, confirming reduced prediction error and stable convergence **(Supplementary Figure 7a, b; Supplementary Table 5)**. Overall, the Aggregator benchmarking suite conclusively establishes CDI as a statistically distinct and top-ranked molecular embedding across all classification and regression tasks. CDI consistently achieved the highest mean performance with the narrowest 95% confidence intervals, demonstrating superior generalization and minimal variance across models and datasets. Comprehensive statistical testing encompassing Mann-Whitney U, ANOVA, and Tukey’s post-hoc analyses confirmed that CDI’s performance advantage is both significant (FDR-corrected, pD<D0.001) and intrinsic to its representational quality, independent of model architecture or task type.

### CDI Establishes Itself as a Top-Tier General-Purpose Molecular Featurizer

To contextualize CDI within the ecosystem of modern molecular representations, we conducted a head-to-head benchmark against the six state-of-the-art Featurizers from which it is derived: ChemBERTa (language-based), GROVER (graph-based), ImageMol (image-based), Signaturizer (bioactivity-based), MOPAC (quantum-mechanical), and Mordred (physicochemical descriptors). The objective was to determine if multimodal fusion could produce an embedding that matches or exceeds the performance of specialized, single-modality experts. Using the extensive benchmark suite of 23 classification datasets (171 tasks), all compounds were encoded using each Featurizer and evaluated through a uniform pipeline of nine machine learning models **(Figure 3a)**. Across this large-scale comparison, CDI emerged as a top-tier, general-purpose molecular representation, consistently matching or surpassing specialized single-modality Featurizers. The results position CDI as a top-tier general-purpose Featurizer. In Featurizer performance, revealed that CDI and Signaturizer formed a clear top-performance cluster, with CDI outperforming descriptor-based (Mordred, MOPAC), graph-based (GROVER), and image-based (ImageMol) methods across both AUC-ROC and Balanced Accuracy, while consistently outperforming all other constituent Featurizers **(Figure 3b)**. Raincloud plots of AUC-ROC and Balanced Accuracy distributions (Mann–Whitney U test, Bonferroni FDR-corrected p < 0.001) show CDI ranking among the top-performing Featurizers, with its performance being significantly superior (p < 0.001) to traditional descriptor-based methods such as MOPAC and Mordred **(Figure 3b, upper)**.

**Figure 3:**
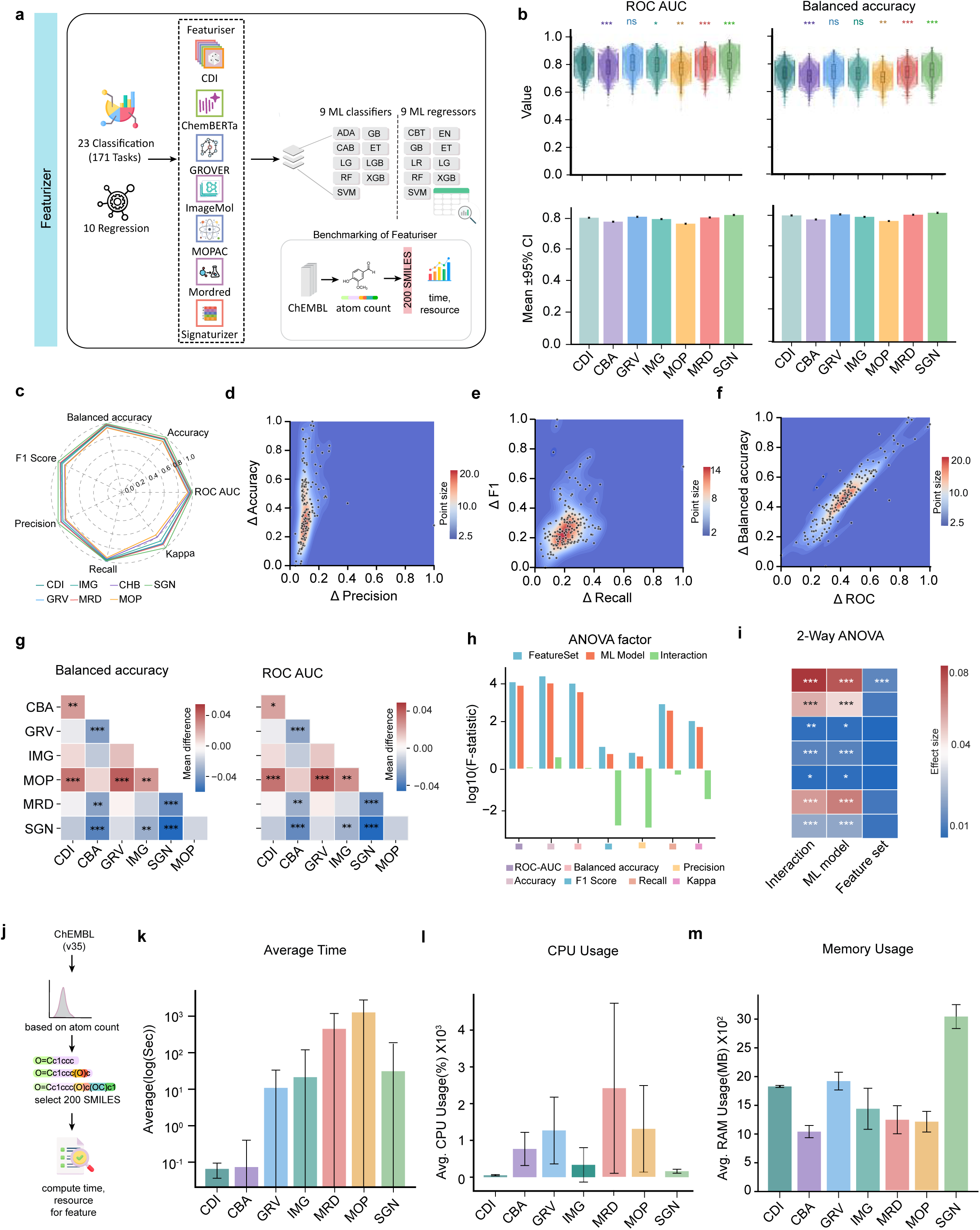
Benchmarking CDI against state-of-the-art molecular Featurizers. **(a)** Schematic of the Featurizer benchmarking strategy. All molecules from the 23 classification datasets (171 tasks) are encoded using CDI and six state-of-the-art Featurizers (ChemBERTa, GROVER, ImageMol, Signaturizer, MOPAC, and Mordred), then evaluated through a uniform pipeline of nine machine learning models. **(b)** Violin plot (above) with distribution of mean AUC-ROC and Balanced Accuracy for all Featurizers across all tasks. Bar plots (below) show Mean performance with 95% confidence intervals for AUC-ROC and Balanced Accuracy for all Featurizers. **(c)** Radar plot synthesizing the performance profiles of all Featurizers across seven key metrics (Balanced accuracy, Accuracy, AUC-ROC, Cohen’s Kappa, Recall, Precision, F1 Score). **(d)** Contour jitter plot of ΔAccuracy vs ΔPrecision confirming CDI’s stronger precision-recall balance between CDI and the best-performing baseline, illustrating CDI’s consistent precision advantage across tasks. **(e)** Contour jitter plot for Δ F1-Score (y-axis) versus Δ Recall (x-axis) confirming CDI’s enhanced balance between sensitivity and predictive reliability. **(f)** Contour jitter plot for Δ Balanced Accuracy (y-axis) versus Δ AUC-ROC (x-axis) demonstrating CDI’s statistical parity or superiority relative to the strongest unimodal Featurizers across evaluation metrics. In panels d-f, points clustered near the origin (0,0) indicate superior or highly competitive performance by CDI. **(g)** One way ANOVA heatmaps with pairwise Tukey’s HSD post-hoc comparison for Balanced Accuracy and AUC-ROC showing significant mean differences (***p < 0.001, ****p < 0.0001) and validating CDI’s statistically distinct performance distribution. **(h)** Bar plot of the log_10_ (F-statistic) from a one-way ANOVA for the factors of FeatureSet, ML Model, and their Interaction, identifying the feature set as the dominant factor. One-way ANOVA across performance metrics (AUC-ROC, Balanced Accuracy, Precision, Recall, F1, Accuracy, Cohen’s Kappa) identifying the feature representation as the primary source of variance across models **(i)** Two-way ANOVA effect-size heatmap (partial η²) demonstrating that the feature-set factor exerts the strongest and most significant influence across all metrics, reaffirming that CDI’s representational quality predominantly dictates predictive success**. (j)** Workflow schematic for computational benchmarking, showing selection of 200 representative SMILES from ChEMBL (v35) based on atom-count distribution and subsequent evaluation of featurization time, CPU, and memory utilization across all methods. **(k)** Bar plots of average feature-generation time (log-scale, seconds) demonstrating that CDI and ChemBERTa are the fastest Featurizers, with total runtimes <1 s per molecule, while descriptor-heavy methods (Signaturizer, Mordred) are considerably slower**. (l)** CPU utilization comparison (percentage ×10³) revealing CDI’s minimal computational load relative to other deep learning and physics-based Featurizers, confirming efficiency and scalability**. (m)** Average RAM consumption (MB ×10^2^) showing CDI’s low and stable memory footprint compared to the higher usage observed for Signaturizer and GROVER, establishing CDI as both performance- and resource-optimal for large-scale molecular screening.

These correspond directly to **Supplementary Figure 8a**, which expands the violin-plot comparison to all additional classification metrics. Accuracy, Kappa, Precision, Recall, and F1-score demonstrate that CDI consistently exhibits the tightest, high-scoring distributions across every metric (Bonferroni-FDR-corrected Mann-Whitney U, p < 0.0001). Bar plots with standard error of mean confirmed that CDI exhibited the highest mean and narrowest CI values, evidencing strong reproducibility and generalization across datasets **(Figure 3b, down)** are paralleled by **Supplementary Figure 8b**, which reports the same confidence-interval analysis for the extended metric set. Across all measures, CDI maintains both the highest mean values and the narrowest confidence intervals, underscoring its exceptional stability and reproducibility over multiple datasets and model architectures. The radar chart **(Figure 3c)** summarizes CDI’s multi-metric dominance, showing that its unified embedding consistently achieved superior values across seven performance indicators: Accuracy, Precision, Recall, F1 Score, Kappa, Balanced Accuracy, and AUC-ROC compared to its constituent Featurizers. CDI formed the outermost and most symmetrical polygon, demonstrating that its latent representation space supports balanced and uniformly strong performance across all evaluation criteria. The critical advantage of the CDI embedding, however, is revealed in a fine-grained analysis of performance differentials. Contour jitter plots of the performance delta (Δ) between CDI and the next-best Featurizer ("Winner") show a consistent and marked superiority for CDI in metrics critical to real-world applications. CDI shows a strong positive skew in Δ Accuracy versus Δ Precision **(Figure 3d)** and Δ F1-Score versus Δ Recall **(Figure 3e)**, indicating it more reliably achieves an optimal balance between identifying true positives and minimizing false positives/negatives across diverse tasks. Furthermore, the tight clustering near the origin for Δ Balanced Accuracy versus Δ AUC-ROC **(Figure 3f)** confirms that CDI consistently outperforms or is at parity with the best single-modality method in overall discriminatory power. This demonstrates that while CDI matches the top-level performance of Signaturizer, it provides a more robust and balanced predictive profile across a broader set of metrics. This comprehensive superiority is further synthesized in that CDI forms the outermost and most balanced profile across seven key metrics **(Figure 3g; Supplementary Table 6)**. Finally, the pairwise ANOVA heatmaps in **(Figure 3g)**, which show mean differences in AUC-ROC and Balanced Accuracy between CDI, are complemented by **Supplementary Figure 8c**. It generalizes statistical comparison to the full set of metrics (Accuracy, Kappa, Precision, Recall, and F1-score), presenting the corresponding mean-difference heatmaps with Bonferroni-FDR–adjusted significance. This statistical analysis was confirmed by a multi-factorial ANOVA. A two-way ANOVA quantified the contribution of the feature set, machine learning model, and their interaction to performance variance, identifying the feature set as the dominant factor across all metrics **(Figure 3h)**. The corresponding effect size heatmap confirms that the feature set has the strongest and most significant effects (p < 0.001), reaffirming that molecular representation quality is the major source of performance variance **(Figure 3i, Supplementary Figure 8)** provides a detailed comparative evaluation, showing that CDI demonstrates strong and consistent results across Accuracy, Cohen’s Kappa, Precision, Recall, and F1 Score, with higher median values and narrow confidence intervals, indicating low variance and excellent reproducibility. A detailed comparative evaluation across multiple classification metrics **(Supplementary Figures 8a-c)** and model-specific analyses **(Supplementary Figures 9a-c, 10a-b)** reveals that CDI consistently performs within the top statistical tier of Featurizers, maintaining high median values, narrow confidence intervals, and minimal variance. Although not always the single best performer in every task, CDI never exhibits statistically significant underperformance relative to leading methods such as Signaturizer or ChemBERTa, demonstrating robust parity and balanced predictive behavior across Accuracy, Kappa, Precision, Recall, F1, and AUC-ROC metrics. This trend is visually reinforced in the contour density plots **(Figures 3d-f)**, where the Δ distributions cluster tightly around the origin, indicating that CDI’s results are generally within the same performance band as the top-performing Featurizer rather than trailing it. Model-wise performance distributions in **(Supplementary Figure 9a-c, 10a-b)** further substantiate this observation. While only a few pairwise comparisons reached statistical significance (p < 0.001), most others remained non-significant (ns), confirming that CDI’s performance margin relative to competitors is narrow but consistent. This pattern suggests a favorable accuracy-stability trade-off, where CDI matches the strongest Featurizers in mean predictive power while exhibiting lower dispersion and higher reproducibility across datasets and algorithms, further confirming CDI’s stable, high-tier performance and robustness across diverse learning frameworks and datasets. Beyond predictive accuracy, CDI achieves competitive computational efficiency, as illustrated in **(Figures 3j-m)**. Using a curated subset of 200 SMILES randomly selected from the ChEMBL (v35) database and stratified by atom count and density distribution of atom count with summary stats of those SMILES **(Figure 3j, Supplementary 10c-d),** we benchmarked average feature extraction time, CPU utilization, and memory footprint. The SMILES were chosen to represent molecular diversity, comprising 100 compounds below the 10th percentile (smaller molecules) and 100 above the 90th percentile (larger, more complex structures) to ensure representative computational scaling. As shown in **(Figures 3k-m)**, CDI ranks among the top three most efficient Featurizers, achieving sub-second average runtime, minimal CPU overhead, and moderate RAM usage. When the Featurizer efficiency and performance metrics are viewed together (**Supplementary Figure 11a)**, for classification datasets, CDI maintains one of the top-three overall ranks across nine evaluation metrics, averaged over multiple datasets. The heatmap visualization demonstrates that CDI’s feature-performance ratio is optimized: while its absolute accuracy is within a narrow margin of the best-performing Featurizer, its substantially lower computational cost provides a superior performance-to-resource balance. In summary, this benchmark establishes that the CDI framework produces a unified embedding that is not only competitive with the best specialized Featurizers but also offers a more balanced and reliable predictive profile. This makes CDI a compelling and versatile choice for a wide range of molecular prediction tasks. Furthermore, CDI in regression tasks consistently achieves decent RMSE **(Supplementary Figure 11b)**. The Featurizer rankings with 95% confidence intervals are summarized in **Supplementary Table 7**.

To assess computational efficiency across different molecular featurization methods, a benchmark was conducted using 200 representative SMILES randomly selected from the ChEMBL (v35) database, stratified by atom count. The average computation time, CPU usage, and memory consumption were recorded for each method: CDI, ChEMBERTa, Grover, ImageMol, Signaturizer, Mordred, and MOPAC. Among the tested approaches, CDI and ChemBERTa demonstrated the fastest feature extraction times, completing processing in less than one second on average, whereas Signaturizer and Mordred required substantially longer runtimes, reflecting higher algorithmic complexity. CPU usage followed a comparable trend, with CDI exhibiting minimal load and Mordred consuming the highest percentage of processing resources. In terms of memory utilization, SGN showed the largest RAM footprint (∼3×10³ M) **(Supplementary Table 8)**, while CDI and ChemBERTa maintained relatively low and stable memory demands. Overall, CDI achieved the most favorable balance between computational speed, CPU efficiency, and memory usage, indicating its strong potential for scalable cheminformatics and bioinformatics applications. To assess CDI’s generalization beyond classification, we extended benchmarking across 10 regression datasets spanning physicochemical and pharmacokinetic properties. The heatmap of regression performance metrics **(Supplementary Figure 11b)** summarizes CDI’s comparative behavior across models, revealing that CDI maintains consistently low error values and stable convergence patterns relative to all other Featurizers. CDI achieves comparable or lower RMSE values than the leading regressors, confirming its robustness in continuous prediction tasks. Across all regressors, Support Vector, Random Forest, Gradient Boosting, XGBoost, CatBoost, and ElasticNet CDI demonstrate reduced Mean Absolute Error (MAE) and Mean Squared Error (MSE) **(Supplementary Figures 12b and 12d, respectively),** indicating a well-calibrated regression space with minimal residual dispersion. The bar plots of median R² scores and test RMSE values **(Supplementary Figure 12c)** further reinforce CDI’s balanced predictive capacity: while it may not exhibit extreme outperformance, CDI consistently delivers top-tier R² values with narrow interquartile ranges, confirming stable model fitting and minimal overfitting. Overall, CDI demonstrates low-error, high-consistency regression behavior, ranking within the top three Featurizers across all continuous prediction metrics. This pattern of low RMSE, narrow MAE/MSE spread, and high R² reproducibility establishes CDI as a statistically stable and computationally efficient general-purpose embedding for both classification and regression tasks (Figures 3g-m; Supplementary Figures 11-12).

### CDI Embodies Superior Chemical Awareness by Discriminating Subtle Molecular Variations

The ability of a molecular representation to distinguish nuanced structural changes is a critical test of its chemical awareness and robustness. We developed the "Complexiser" module to quantitatively evaluate whether CDI’s multimodal fusion results in a more chemically sensitive embedding compared to single-modality Featurizers. We tasked all models with separating two types of subtle molecular variations: chiral enantiomers versus non-chiral molecules and kekulized versus their non-kekulized forms, which explicitly define aromatic bond alternation **(Figure 4a)**. For the chiral separation task, using a balanced dataset of 1,000 chiral and 1,000 non-chiral molecules **(Figure 4b)**, CDI demonstrated superior latent structure organization. Elbow curve analysis of the PCA-transformed features showed that CDI’s embedding space facilitated a more distinct clustering, as indicated by a more pronounced drop in within-cluster inertia **(Figure 4c)**. Visualization of the first two principal components confirmed this quantitatively; only CDI and ChemBERTa achieved clear visual separation between chiral and non-chiral clusters, while all other Featurizers showed significant overlap **(Figure 4d)**. The t-SNE projections for this task, as shown in **Supplementary Figure 13a**, reveal minimal visual separation across all Featurizers, as indicated by very low silhouette scores. Quantitative clustering metrics comparing t-SNE and PCA embeddings confirm that PCA consistently yields higher clustering quality for capturing stereochemical distinctions **(Supplementary Figure 13b)**. These comparisons demonstrate that PCA projections more effectively capture stereochemical variance than t-SNE for this dataset. The kekulization task provided an even more rigorous test for features to electronic structure representations in a balanced dataset of 1,522 non-kekulized and 1,522 kekulized SMILES **(Figure 4e)**. Here, CDI’s multimodal advantage became overwhelmingly clear. The elbow curve for CDI indicated the most well-defined cluster separation among all Featurizers **(Figure 4f)**. Crucially, in the PCA projection, only the CDI embedding achieved a near-complete separation of the two forms **(Figure 4g).** ChemBERTa, the second-best performer on the chiral task, showed substantial overlap, while all other methods failed entirely to distinguish the kekulized states. The t-SNE scatter plots for the kekulized dataset in **Supplementary Figure 13c** exhibit clear and well-defined clustering, with CDI and ChemBERTa showing the most distinct separations. The comparison of clustering performance metrics in **Supplementary Figure 13d** shows that both PCA and t-SNE achieve improved metrics for the kekulized data compared to the chiral case, with PCA again outperforming t-SNE overall. Additionally, CDI achieves among the highest silhouette and Calinski-Harabasz scores. This indicates that CDI’s integration of quantum (MOPAC) and graph-based (GROVER) features provides a unique sensitivity to electronic and bond-level nuances that are lost in language-based or descriptor-based approaches. These results provide compelling evidence that the CDI framework learns a chemically intelligent representation. Its ability to consistently and clearly segregate both stereochemical and electronic molecular variants establishes that its multimodal fusion process captures a deeper and more fundamental level of chemical structure than any single constituent Featurizer. The Complexiser analysis thus establishes CDI as a chemically intelligent representation capable of differentiating molecular variants across both configurational and electronic dimensions. Cross-language validation confirmed near-perfect embedding equivalence between the Python and R implementations (mean cosine similarity ∼0.99) **(Supplementary Figure 14h)**, confirming CDI’s full interoperability and reproducibility as a unified featurization platform for large-scale cheminformatics and bioinformatics applications.

**Figure 4:**
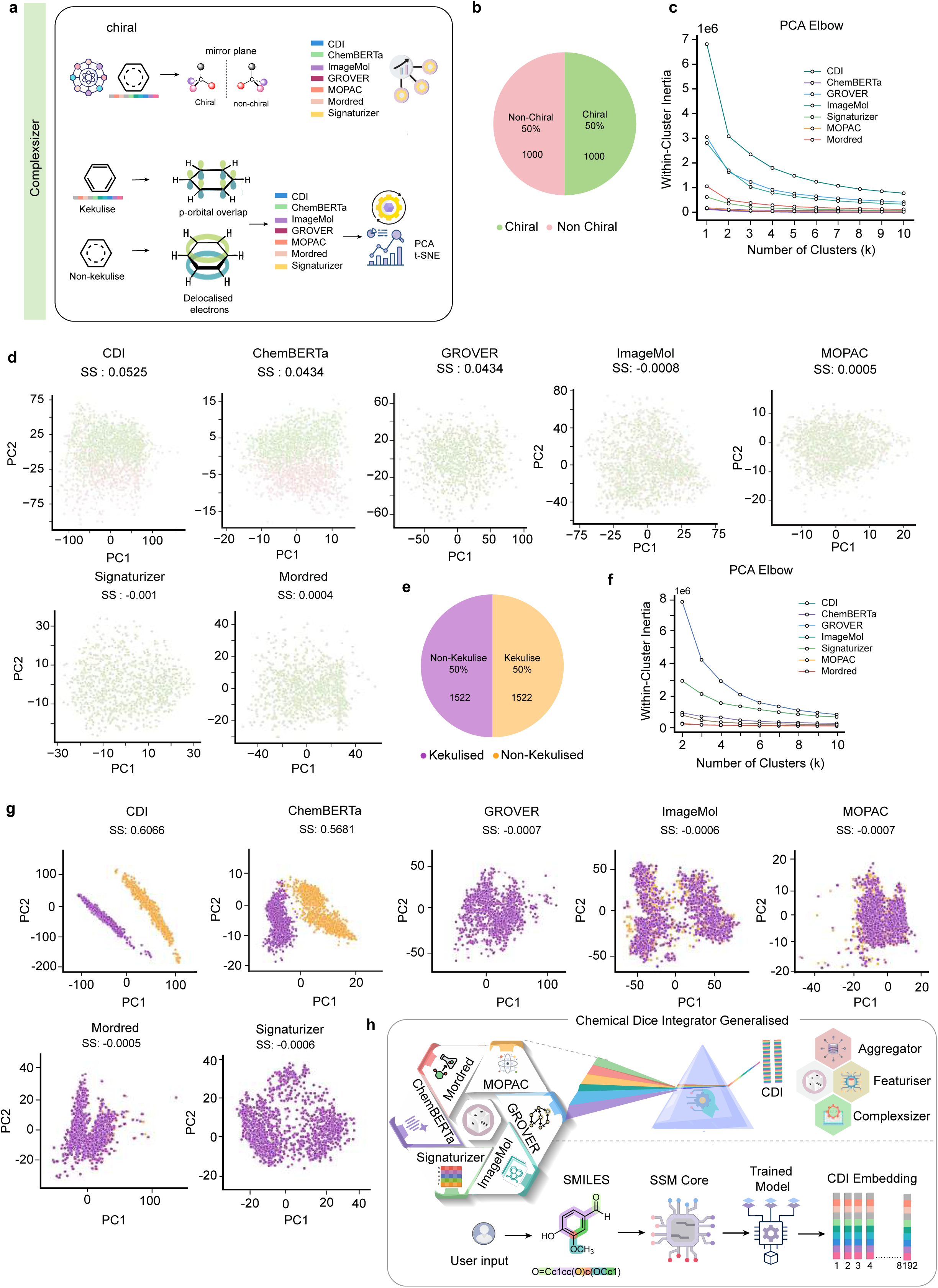
The Complexiser analysis demonstrates CDI’s superior sensitivity to nuanced chemical structures. **(a)** Schematic overview of the Complexiser experimental design, comparing the ability of seven Featurizers CDI, ChemBERTa, GROVER, ImageMol, MOPAC, Signaturizer, and Mordred to distinguish chiral versus non-chiral molecules and kekulized versus non kekulized SMILES forms. **(b)** Pie chart showing the balanced dataset of the chiral versus non-chiral (n=1,000 molecules each) curated for stereochemical discrimination analysis. **(c)** Elbow curve analysis (within-cluster Inertia vs. number of clusters, k) derived from PCA embeddings, for all Featurizers on the chiral dataset, with CDI showing a pronounced curve indicative of well-defined cluster structure. **(d)** PCA projections (PC1 vs. PC2) of the chiral/non-chiral dataset for all Featurizers, revealing that only CDI and ChemBERTa achieve clear visual separation between the two classes. **(e)** Pie chart showing the balanced distribution of the kekulized versus non-kekulized SMILES (n=1,522 molecules each) prepared for structure sensitivity assessment. **(f)** Elbow-curve plots of within-cluster inertia for the kekulized dataset confirming CDI’s most distinct elbow and strongest separation, indicative of enhanced awareness of aromatic bond alternation. **(g)** PCA projections (PC1 vs PC2) for the kekulized versus non-kekulized dataset, demonstrating that CDI alone achieves near-complete separation of kekulized and non-kekulized forms, underscoring its unique sensitivity to bond-order information. **(h)** Graphical abstract summarizing the Chemical Dice Integrator (CDI) framework. The figure illustrates the multimodal integration of six orthogonal representations MOPAC (quantum), Signaturizer (bioactivity), ChemBERTa (language), ImageMol (image), Mordred (physicochemical), and GROVER (graph), within the CDI-Basic autoencoder, followed by sequence-to-embedding translation via the Mamba State-Space Model (SSM) core. The unified CDI embedding supports three operational modules: Aggregator (benchmarking and statistical evaluation), Featurizer (performance evaluation), and Complexiser (structural sensitivity), collectively enabling chemically aware, scalable, and reproducible molecular representation learning.

## Discussion

The rapid development of specialized molecular Featurizers, from graph networks (GROVER ^4^) to language models (ChemBERTa ^5^), has paradoxically created a fundamental bottleneck in molecular machine learning. Researchers are often forced to make a priori choices about molecular representation, locking models into a single, potentially suboptimal "view" of chemical space and limiting their generalizability across diverse prediction tasks ^13^. Our work introduces the Chemical Dice Integrator (CDI) to resolve this fragmentation. We demonstrate that a purposefully designed, hierarchical fusion of six complementary molecular modalities yields a unified embedding that is not only more predictive and robust than any single Featurizer but also uniquely sensitive to subtle structural nuances, all while being computationally scalable for real-world deployment. The principal strength of CDI lies in its ability to generate synergistic representation learning. This is conclusively evidenced in the Aggregator benchmark, where CDI’s performance consistently surpassed a wide array of standard dimensionality reduction techniques, including linear methods like PCA and non-linear manifold learners like t-SNE and Isomap. This performance gap suggests that CDI’s advantage originates from its two-tiered autoencoder architecture, which actively learns the complex, non-linear relationships that interconnect different chemical domains, such as correlating quantum mechanical features with bioactivity profiles, rather than merely compressing the feature space ^12^. This result strongly supports the growing consensus that the next frontier in molecular ML lies in intelligent integration, moving beyond competitions between isolated, specialized models ^14^. As a direct Featurizer, CDI establishes itself as a top-tier, general-purpose tool. It matched the powerful, bioactivity-driven Signaturizer ^9^ in overall discriminatory power (AUC-ROC) but achieved a statistically significant advantage in balanced metrics, such as F1-score and precision. This indicates that CDI delivers a more reliable and robust predictive profile across chemically diverse and often imbalanced datasets, a critical asset for practical discovery pipelines where the cost of false positives and negatives is high.

Beyond predictive performance, CDI introduces pivotal computational advancements. The CDI-Generalised model, which leverages the efficient Mamba State-Space Model (SSM) ^15^, addresses a key scalability problem. It provides a direct, high-speed pathway from a SMILES string to the unified embedding, decoupling the rich, multimodal representation from the computationally expensive process of generating all six source features during inference. This enables the breadth of an ensemble with the latency of a single model. Furthermore, the fully containerized API deployment transforms CDI into a production-ready molecular intelligence engine, ensuring reproducibility and hardware flexibility ^16^ and directly addressing the operational gap in deploying complex AI within chemical workflows.

Perhaps the most profound validation of CDI’s chemical intelligence comes from the Complexiser analysis. CDI’s superior ability to distinguish between kekulized and canonical SMILES forms, a task that probes sensitivity to electronic structure, demonstrates that its embedding space captures the nuanced "grammar" of chemical bonding, a feat that eluded most single-domain Featurizers. We acknowledge that the current framework requires an upfront pre-training cost for the fusion engine and is inherently constrained by the scope of its six constituent Featurizers. Future work will focus on incorporating 3D structural and dynamical information to model target-specific interactions more directly ^17^. In conclusion, the CDI framework successfully bridges the critical gap between the power of multimodal chemical intelligence and the practical demands of computational efficiency and deployability. It provides a unified, chemically aware, and scalable foundation poised to accelerate discovery in drug design and materials science. Future work shall be focused on extending CDI with 3D-aware conformer embeddings (e.g., SchNet or UniMol) to capture spatial molecular dynamics, exploring multitask and few-shot transfer learning paradigms to enhance generalizability across chemical domains, and ensuring full FAIR compliance so that CDI remains findable, accessible, interoperable, and reusable for the broader research community. In summary, CDI bridges the long-standing divide between representational power and computational efficiency, providing a unified, chemically aware, and scalable foundation for next-generation discovery pipelines. Its multimodal fusion design, combined with deployable inference and electronic-structure awareness, positions CDI as a cornerstone for integrating chemical intelligence into drug discovery, materials design, and beyond.

## MATERIAL AND METHODS

### Architecture of the Chemical Dice Integrator (CDI) Generalised Framework

The Chemical Dice Integrator (CDI) is a hierarchical, multimodal architecture designed to unify diverse molecular representations into a single, high-information, generalised embedding. The framework comprises two core components: (1) CDI-Basic: an autoencoder-driven multimodal fusion engine that generates a unified embedding, and (2) CDI-Generalised: a Mamba State-Space Model (SSM) based regression head that learns to map raw molecular sequences directly to this embedding space.

#### Multimodal Fusion (CDI-Basic)

The workflow initiates with curated SMILES strings, sourced from ChEMBL (v35) database. Each molecule is transformed into six complementary embedding modalities to capture orthogonal biochemical and structural information: Mordred (physicochemical descriptors), GROVER (graph-based topology), ImageMol (structural image-based features), Signaturizer (bioactivity profiles), MOPAC (quantum mechanical properties), and ChemBERTa (language model-based semantics). For each molecule *x^i^*, these embeddings form a holistic feature set, denoted as

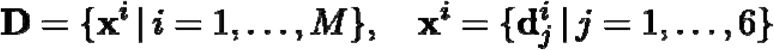

where each M is the total number of molecules, and 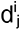 for j = 1 to 6 represents j^th^ semantic embedding. To integrate these modalities, CDI employs a two-tiered autoencoder strategy for hierarchical semantic fusion. In the first stage, six dedicated Semantic Commonality Autoencoders (SCA) learn inter-modality dependencies. Each SCA_j_ is trained to reconstruct j^th^ embedding from concatenated latent information of the other five. This process encodes the input into a latent representation 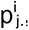, which is dimensionality-matched to the target embedding. The training objective for each SCA combines a mean squared error (MSE) loss (L_MSE_) with an auxiliary reconstruction error (RE) loss term to ensure close alignment between latent representation and target modality. For every embedding 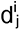, model builds one SCA block, with input to that SCA block being 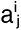, where

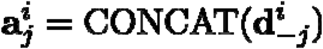

The SCA encoder compresses this concatenated vector into a latent representation 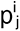

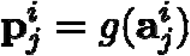

where.

Each SCA decoder f(.) reconstructs 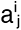 from 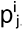 Along with using the traditional MSE loss to train the encoder and decoder blocks, we introduce an additional loss term to specifically train the encoder block using the Reconstruction Error (RE), which is calculated between 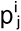 and 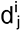 to maintain structural coherence across modalities. Each SCA is trained with this dual-loss objective combining Mean Squared Error (MSE) and Reconstruction Error (RE):

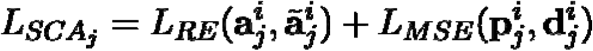

Where L_MSE_ and L_RE_

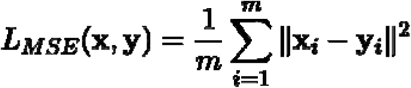

where x, y i ii

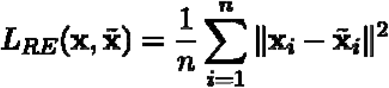

where x, 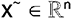

The ReLU activation ensures non-linearity and sparsity during training

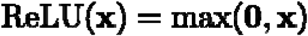

Each SCA captures how modalities relate to one another, focusing on preserving the unique properties of the modality left out during training. The first term ensures input reconstruction. The second term enforces alignment between the latent embedding p^i^ and the target embedding d^i^ . The MSE term drives the model to learn the structural patterns shared among the five embeddings, while the reconstruction error (RE) teaches each SCA how these five modalities can be combined to reconstruct the sixth, omitted one. Together, these objectives enable the model to preserve the unique characteristics of each modality while also capturing its relationships with the remaining modalities. In this way, each SCA learns the semantic information common to all modalities except the one it aims to predict. The six information-rich, intermediate, latent representations are concatenated and passed into the Super-Embedding Autoencoder (SEA) to produce a unified representation, which we call the ‘Super Embedding’.

Encoder: compresses the joint information into a single super-embedding. Decoder: reconstructs concatenated latent vector. The SEA compresses the combined information into a unified latent space and reconstructs the concatenated latent input. The decoder tries to reconstruct either the concatenated latent vector (local consistency) or the original embeddings (global reconstruction). Losses from all SCA blocks and the super-autoencoder are combined for joint optimization. The overall objective minimizes total reconstruction. The total hierarchical training objective jointly optimizes all local (SCA) and global (SEA) reconstruction losses:

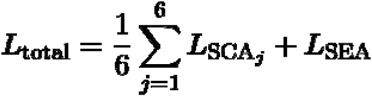

After convergence, reconstruction quality was quantified using MSE and RE, and latent vector similarity. The SEA further expands this integrated semantic space to yield the final CDI super-embedding:

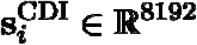

Training drives each encoder–decoder pair to preserve semantic relationships while removing redundant and retaining useful information. The SEA is trained to compress this concatenated input into a single, high-dimensional CDI super-embedding (8192 dimensions) that encapsulates the joint semantics of all modalities, minimizing the reconstruction error to ensure a cohesive integration.

### Sequence-to-Embedding Mapping (CDI-Generalised)

The CDI-Generalised model learns to map raw SMILES sequences directly to the CDI-Basic embedding space using a Mamba SSM core. Subsequently, the aforementioned fused CDI embeddings serve as the training targets for the Mamba SSM mapping network. This component enables direct inference from SMILES sequences. The SMILES strings are first tokenized and passed through an input embedding layer. The Mamba SSM ^15^ core then processes these sequences, efficiently capturing both short and long-range dependencies via its state-space dynamics. The final hidden state is fed through a regression head to predict an embedding vector that approximates the target CDI (8192) embedding vectors. This model is optimized using a composite loss function that combines MSE, thereby preserving both the magnitude and angular relationships within the embedding space. The standard training loop involves a forward pass, loss computation, gradient backpropagation, and parameter updates using the AdamW optimizer. Upon convergence, the model provides a direct, high-fidelity mapping from raw SMILES strings to the high-dimensional CDI representation. The final hidden state was passed through a regression head to predict an embedding vector approximating the target CDI representation. The SSM was optimized with a MSE Loss that preserves both magnitude and angular relationships:

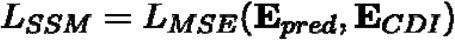

In summary, the CDI-Generalised architecture achieves multimodal unification through hierarchical autoencoder fusion and robust sequence-based inference via the Mamba SSM. It effectively bridges six distinct chemical feature spaces and enables the generation of information-rich, unified embeddings directly from molecular structures, facilitating scalable and modality-independent chemical representation learning. The CDI-Basic architecture comprised approximately 1.09 billion trainable parameters, indicating a high-capacity network designed to capture complex, non-linear feature relationships. The model was optimized using SGD with a learning rate of 0.05, a configuration selected to balance convergence speed and stability. The Mean Squared Error (MSE) loss function was employed to minimize reconstruction error between the input and output, descriptor and latent-space embeddings, ensuring effective feature compression. Training was performed for 600 epochs using the entire dataset without a separate validation partition, thereby allowing for the full utilization of available samples and enabling representation learning within the high-dimensional feature space. CDI-Generalised training were performed using random seed 42; 90:10 train/validation trained with AdamW (lr=1e-5, weight decay=1e-2), batch size=32, and training was conducted for 30 epochs.

### API and Deployment Architecture

To ensure scalable and reproducible deployment, the CDI-Generalised framework was implemented as a containerized web service. The core logic, comprising the multimodal autoencoders (CDI-Basic), the Mamba SSM model (CDI-Generalised), and data preprocessing units, was modularized into a FastAPI application. Discrete RESTful endpoints were created for critical tasks: SMILES validation, tokenization, embedding generation, and inference computation. Outputs were serialized into standardized CSV formats. The entire application was containerized using Docker, encapsulating all dependencies (PyTorch, RDKit, Mamba SSM) to guarantee consistency across different hardware and operating systems. The API server was built with Uvicorn workers and fronted by an NGINX load balancer to efficiently manage high-throughput, concurrent requests. The inference layer was designed to utilize GPU (CUDA), optimizing for both speed and cost efficiency. System resource usage (RAM, CPU, I/O) was continuously logged for monitoring and dynamic resource management. This architecture enables seamless integration into larger computational workflows and deployment on container orchestration platforms, such as Kubernetes, for cloud-scale production use. Docker images and configuration files are provided in the Zenodo for reproducibility.

### Dataset Curation for Test Architecture of CDI

To rigorously evaluate the CDI framework, we curated a comprehensive suite of benchmark datasets spanning diverse prediction tasks. All datasets were standardized into canonical SMILES format and annotated with their respective experimental activity or property values. We assembled 23 datasets totaling 171 distinct classification tasks comprising roughly 0.92 million unique molecules **(Supplementary Table 2)**. These covered key areas in toxicology (AMES, Drug-Induced Liver Injury (DILI), Side Effect Resource (SIDER) ^18^, Toxicology in the 21st Century (Tox21), Toxicity Forecaster (ToxCast)), ADMET properties (PGP efflux, Blood-brain barrier (BBB) permeability, CYP450 inhibition for 2C9, 2D6, and 3A4 isoforms, hERG blockage, bioavailability) ^19^, and bioactivity (HIV, beta-secretase (BACE), CAV3, PCBA) ^20^. We also selected 10 datasets representing continuous molecular properties, including Caco-2 permeability, lipophilicity, solubility, hydration free energy, plasma protein binding rate (PPBR), volume of distribution (Vdss), half-life, clearance, BACE binding affinity, and LD50 Regression **(Supplementary Table 3)**. Datasets were sourced from public repositories such as ChEMBL ^21^, ToxCast ^22^, and the Therapeutic Data Commons (TDC) to ensure non-redundant, experimentally validated entries. This collection provides broad coverage from focused, small-scale assays to large-scale screening panels, enabling a robust assessment of model generalizability across the chemical and pharmacological space.

### Performance Benchmarking: Aggregator Module

The Aggregator module was designed to systematically compare the predictive performance of CDI embeddings against established feature-space transformation techniques. For each of the curated benchmark datasets, molecular structures were encoded using CDI and eight other methods: PCA (Principal Component Analysis), kPCA (kernel Principal Component Analysis), ICA (Independent Component Separation), CCA (Canonical Correlation Analysis), Isomap (Isometric Mapping), LLE (Locally Linear Embedding), t-SNE (t-Distributed Stochastic Neighbor Embedding), and RKS (Randomized kernel Approximation). Each resulting feature set was evaluated using a uniform machine-learning pipeline. We employed nine models: Support Vector Machine (SVM), Logistic Regression (LR), Random Forest (RF), Gradient Boosting Classifier (GBC), Gaussian Naïve Bayes (GNB), K-Nearest Neighbors (KNN), Extra Trees (ET), Stochastic Gradient Descent (SGD), and Multi-Layer Perceptron (MLP). For regression tasks, analogous regressors were used, including Extreme Gradient Boosting (XGBoost). All models were trained under identical conditions, including hyperparameter tuning via cross-validation, to ensure that performance differences were attributable solely to the quality of the embeddings. Performance was assessed using the AUC-ROC, Balanced Accuracy, F1 Score, Recall, Cohen’s Kappa, precision, R², and MAE, MSE, and RSME for classification and regression, respectively. Performance distributions were compared pairwise using Mann-Whitney U tests with Bonferroni-FDR correction. Ninety-five-percent confidence intervals were estimated by bootstrapping (n = 1000 resamples). To quantify variance contributions, a two-way ANOVA was applied with feature representation and machine-learning model as fixed factors and dataset identity as a random effect. The ANOVA model generated F-statistics and p-values for main effects and interaction terms. Partial η² values were computed to estimate the effect size of each factor, providing a quantitative measure of variance attribution. In addition, a one-way ANOVA followed by Tukey’s Honest Significant Difference (HSD) post-hoc test with Bonferroni-FDR correction was used to assess pairwise differences among all feature representations. Mean rank aggregation was performed across all metrics and models to generate the comparative ranking matrices.

### Performance Benchmarking: Featurizer Module

The Featurizer module was implemented to directly benchmark CDI as a molecular representation against six state-of-the-art Featurizers from which it is derived: ChemBERTa (language-based), GROVER (graph-based), ImageMol (image-based), Signaturizer (bioactivity-based), MOPAC (quantum mechanical), and Mordred (physicochemical descriptors). Using the same benchmark datasets and uniform machine-learning pipeline described in the Aggregator module, we compared the performance of these seven Featurizers. The evaluation focused on a suite of seven metrics: AUC-ROC, Balanced Accuracy, F1-score, Precision, Recall, Accuracy, and Cohen’s Kappa to provide a comprehensive view of model behavior, particularly on imbalanced datasets. Statistical significance of performance differences across Featurizers was assessed using a one-way ANOVA, accounting for inter-dataset variability similar to the Aggregator module.

### Computational Efficiency Evaluation of Molecular Featuriser Methods

To systematically assess the computational efficiency of different molecular representation paradigms, we utilized the ChEMBL v35 dataset as a large-scale and chemically diverse source of compounds. We first computed the atom count for all molecules (canonical SMILES) and generated a density distribution to characterize molecular size variation. The distribution exhibited substantial right-skewness, indicating a predominance of small molecules with a long tail of larger compounds. Atom-count distributions were computed for all ChEMBL v35 SMILES. Molecules were sorted in descending order, and 100 samples were drawn from the 10th and 90th percentiles (200 total) to represent small and large molecules. We benchmarked the featurization time, CPU utilization, and GPU usage across a diverse set of molecular representation tools encompassing multiple learning paradigms: CDI, ChemBERTa, GROVER, ImageMol, Signaturizer, MOPAC, and Mordred. Each tool was executed using default or recommended configurations to generate molecular embeddings or feature vectors for the selected 200 molecules. Runtime and hardware utilization metrics were recorded for both CPU-only and GPU-enabled environments, providing a quantitative comparison of computational efficiency across representation modalities. To ensure reproducibility and isolate system-level variability, all featurization pipelines were encapsulated within Docker containers, allowing uniform execution environments across methods. This containerized setup ensured that runtime and hardware utilization metrics were measured under identical conditions for both CPU-only and GPU-enabled configurations, enabling a fair and controlled comparison of computational efficiency across molecular representation approaches. All benchmarks were executed in Docker containers using identical compute environments (CPU : AMD Ryzen Threadripper 3960X 24-Core Processor 24C | 48T, RAM 256GB, GeForce RTX 3090 24 GB, Operating system Ubuntu : 20.04.6 LTS).

### Structural Sensitivity Analysis: Complexiser Module

The Complexiser module was developed to quantitatively evaluate the sensitivity of molecular embeddings to subtle structural changes. We tested the ability of CDI and six baseline Featurizers to distinguish between two types of molecular representations: Chiral vs. Non-Chiral Molecules: A balanced dataset taken from Yoshikai *et al.* ^23^ of 1,000 chiral and 1,000 non-chiral molecules **(Supplementary Table 9)**. Kekulized vs. Non-kekulized SMILES: A balanced dataset of 1,522 molecules from the BACE dataset (Moleculenet) was prepared, and RDKit was used to convert the original non-Kekulized SMILES into their Kekulized forms **(Supplementary Table 10)**. Using this dataset of 3044 molecules (kekulized and non-kekulized), molecular features were computed with various Featurizers, including CDI, Signaturizer, GROVER, ChemBERTa, ImageMol, Mordred, and MOPAC. For each Featurizer and dataset pair, the high-dimensional embeddings were projected into a 2D space using PCA and t-SNE for visualization. The quality of separation between the two classes (chiral/non-chiral) was quantified using cluster validation metrics: the Silhouette Score, Davies-Bouldin Index, and Calinski-Harabasz Score.

### CDI Python package and R library

The Chemical Dice Integrator (CDI) was implemented as a cross-platform molecular featurization framework available in both Python and R, ensuring identical embeddings and reproducibility across environments. The Python package, developed using FastAPI and PyTorch, integrates RDKit for SMILES validation and preprocessing, generating 8192-dimensional embeddings directly from input SMILES via the CDI-Generalised Mamba State-Space Model. The R interface mirrors this functionality through reticulate, enabling seamless access to the same backend for feature generation and validation. Both implementations operate within a unified Dockerized architecture, ensuring consistent dependencies, automatic API uptime, and full compatibility with GPU and CPU hardware. Cross-language validation confirmed embedding equivalence (cosine similarity = 1), establishing CDI as a fully interoperable and reproducible featurization platform for large-scale cheminformatics and bioinformatics applications.

## Statistical Analysis

All statistical analyses were performed using SciPy (v1.15.3) and StatsModels (v0.14.2) Python packages. Pairwise model comparisons were conducted using the Mann-Whitney U test to assess median differences in non-parametric distributions across metrics (AUC-ROC, Balanced Accuracy, Accuracy, Precision, Recall, F1, and Cohen’s κ). *P*-values were adjusted for multiple testing using the Bonferroni False Discovery Rate (FDR) correction. A one-way ANOVA was used to evaluate the effect of feature representation on performance metrics, followed by Tukey’s HSD post-hoc test to identify statistically significant pairwise differences between Featurizers, with multiple-testing correction. A two-way ANOVA quantified variance contributions from feature set, machine learning model, and their interaction, confirming feature representation factor influencing performance across metrics. Statistical significance was set at pL<L0.05, with significance levels denoted as *L<L0.05, **L<L0.01, ***L<L0.001.

## Data and Code Availability

A Python package for CDI is provided via pip: https://test.pypi.org/project/ChemicalDice/ . R library and python package can be accessed via Github (https://github.com/the-ahuja-lab/ChemicalDice). CDI is free for academic institutions; however, a commercial license key is mandatory for commercial usage. The source code of CDI is available on the Zenodo project. All computational resources used, along with their corresponding versions, are listed in Supplementary Table 1. The datasets used in this project can be found at the following links: 2.23 million molecules of ChEMBL (Zenodo: https://zenodo.org/records/17582661), molecular property prediction summary datasets of classification (Supplementary Table 2), and regression (Supplementary Table 3).

## Acknowledgments

The authors thank the IT-HelpDesk team at IIIT-Delhi for their assistance with computational resources. We thank all the members of the Ahuja lab for their intellectual contributions at various stages of this project. The Ahuja lab is supported by the Ramalingaswami Re-entry Fellowship (BT/HRD/35/02/2006), and a research grant (BT/PR52020/AI/133/180/2024) by the Department of Biotechnology, Ministry of Science & Technology, Government of India, and an intramural Start-up grant from Indraprastha Institute of Information Technology-Delhi.

## Author Contributions

The study was conceived and supervised by G.A. S.K. built the entire foundation of ChemicalDice, and M.G. and S.S. designed the architecture of CDI-generalised. S.K.M assisted S.K. in data gathering and machine learning pipeline setup. S.S, S.C, S.D, A.S, V.G, S.A, R.S, S.S, A.K.S, and A.M. performed large-scale validations of CDI on various datasets. D.S. guided the statistical analysis, and N.A.M. helped with computing various descriptors and its integration. G.A, S.S. and S.K. draft the figures and the manuscript. All authors read and approved the manuscript.

## SUPPLEMENTARY FIGURE LEGENDS

Supplementary Figure 1: Mathematical formulation, dataset preparation, and hyperparameter configuration of the Chemical Dice Integrator (CDI) **(a)** Step-wise mathematical formulation of the CDI architecture showing encoder–decoder mappings, reconstruction objectives, and loss functions. The equations define the Semantic Commonality Autoencoder (SCA) and Super-Embedding Autoencoder (SEA) training procedures, integrating mean-squared-error (MSE) and reconstruction-error (RE) components with ReLU activation. The formulation details latent vector concatenation, dimensional alignment across modalities, and the composite optimization objective for hierarchical fusion. **(b)** Dataset curation workflow for CDI development. The ChEMBL v35 corpus (2.4 million molecules) was filtered to 2.23 million after ChemBERTa-based preprocessing and removal of entries failing MOPAC quantum-mechanical computation, yielding a high-quality, non-redundant dataset for model training. **(c)** Table summarizing model configuration and training hyperparameters for CDI-Simple and CDI-Generalised variants. The Generalised model, trained with AdamW optimizer and a smaller batch size, exhibits faster convergence and lower reconstruction loss with substantially fewer parameters (0.108 B vs 1.09 B).

Supplementary Figure 2. Statistical comparison of feature representations across evaluation metrics. **(a)** Boxplots comparing Accuracy, Precision, Recall, F1 Score, and Cohen’s Kappa across all aggregation methods, highlighting CDI’s highest medians and narrowest interquartile ranges. The consistent clustering of CDI values at the upper range demonstrates superior stability and generalization. **(b)** Raincloud plots with violin distributions depicting the performance spread for each method across all benchmark tasks. CDI exhibits the tightest and most elevated distributions for all metrics, reflecting strong cross-dataset robustness and minimal variance. Statistical significance was assessed using the Mann–Whitney U test (***p < 0.001). **(c)** Mean ± 95% CI bar plots confirming CDI’s consistent stability and accuracy for Accuracy, Balanced Accuracy, Recall, F1 Score, Precision and Cohen’s Kappa. **(d)** Pairwise comparison heatmaps using Tukey’s Honest Significant Difference (HSD) post-hoc test for Accuracy, Balanced Accuracy, Recall, F1 Score, Precision, and Cohen’s Kappa. Positive mean differences (red) indicate superior CDI performance, while blue regions represent lower-scoring baselines. Significance thresholds are denoted as *p < 0.05, **p < 0.01, ***p < 0.001, ****p < 0.0001

Supplementary Figure 3. Comparative performance of feature representations across classifiers: Precision, Recall, and AUC-ROC. **(a)** Boxplots illustrating Precision values for nine machine-learning classifiers (AdaBoost, CatBoost, Extra Trees, Gradient Boosting, LightGBM, Logistic Regression, Random Forest, SVM, and XGBoost) across all feature aggregation approaches. CDI consistently exhibits the highest median precision and lowest variance, confirming stable discrimination between active and inactive classes. **(b)** Boxplots depicting the Recall across nine classifiers (AdaBoost, CatBoost, ExtraTrees, GradientBoosting, LightGBM, LogisticRegression, RandomForest, SVM, and XGBoost), with statistical significance assessed using the Mann - Whitney U test showing CDI’s superior sensitivity in identifying true positives relative to all baseline methods. **(c)** Boxplots of AUC-ROC across nine classifiers (AdaBoost, CatBoost, ExtraTrees, GradientBoosting, LightGBM, LogisticRegression, Random Forest, SVM, and XGBoost) with statistical significance assessed using the Mann Whitney U test.

Supplementary Figure 4. Classifier-wise comparison of Cohen’s Kappa, Balanced Accuracy, and Accuracy.**(a)** Boxplots showing the performance of nine classifiers based on Cohen’s Kappa. Kernel methods (KPCA, Isomap) rank next best, whereas stochastic embeddings (t-SNE, RKS) perform least reliably. Statistical significance were assessed using the Mann-Whitney U test. **(b)** Boxplots showing the performance of nine classifiers based on Balanced Accuracy with statistical significance were assessed using the Mann Whitney U test. **(c)** Boxplots showing the performance of nine classifiers based on Accuracy statistical significance were assessed using the Mann Whitney U test.

Supplementary Figure 5. Cross-dataset and cross-model performance heatmaps for classification tasks. **(a)** Heatmaps summarizing classifier performance (AUC-ROC, Accuracy, Balanced Accuracy, F1 Score, Precision, Recall, Cohen’s Kappa) across six benchmark datasets (ToxCast, Tox21, SIDER, BACE, PCBA, TDC). Each cell represents the mean z-score-normalized metric per model–dataset pair. Boosting-based algorithms (LightGBM, XGBoost, and Gradient Boosting) consistently achieve the highest scores when trained on CDI embeddings, confirming strong generalizability across data domains.

Supplementary Figure 6. Cross-dataset regression benchmarking across models. **(a)** Regression heatmaps showing model dataset with nine regressor performances for R², MAE, RMSE, and MSE metrics across nine regressors (Support Vector, Random Forest, Gradient Boosting, Extra Trees, XGBoost, CatBoost, Lasso, and ElasticNet) over ten physicochemical datasets, showing CDI’s uniformly superior correlation (high R²) and lowest errors. **(b)** Bar plots of median R² comparisons across nine regressors (support vector regressor, Random Forest, Gradient boosting, extra trees, XGBoost regressor, CatBoost Regressor, Lasso Regressor, Elasticnet). **(c)** RMSE bar plots across nine regressors (support vector regressor, Random Forest, Gradient boosting, Extra trees, XGBoost regressor, CatBoost Regressor, Lasso Regressor, Elasticnet) with Aggregators (CDI, CCA, ICA, Isomap, KPCA, LLE, PCA, RKS, t-SNE).

Supplementary Figure 7. Regression error analysis across feature representations. **(a)** MSE bar plots across nine regressors (Support vector regressor, Random Forest, Gradient boosting, extra trees, XGBoost regressor, CatBoost Regressor, Lasso Regressor, Elasticnet) with Aggregators (CDI, CCA, ICA, Isomap, KPCA, LLE, PCA, RKS, and tSNE). **(b)** MAE bar plots across nine regressors (Support vector regressor, Random Forest, Gradient boosting, extra trees, XGBoost regressor, CatBoost Regressor, Lasso Regressor, Elasticnet) with Aggregators (CDI, CCA, ICA, Isomap, KPCA, LLE, PCA, RKS, and tSNE).

Supplementary Figure 8. Comparative evaluation of Featurizers across multiple metrics and models. **(a)** Violin and Raincloud with distribution plots of Accuracy, Cohen’s Kappa, Precision, Recall, and F1 showing CDI and other state-of-the-art Featurizers CDI, ChemBERTa (CBA), GROVER (GRV), ImageMol (IMG), Signaturizer (SGN), MOPAC (MOP), and Mordred (MRD). **(b)** Mean ± 95% CI of SEM bar plots for Precision, Recall, Cohen’s Kappa, F1Score, Accuracy for featurizers CDI, ChemBERTa (CBA), GROVER (GRV), ImageMol (IMG), Signaturizer (SGN), MOPAC (MOP), and Mordred (MRD). **(c)** One way ANOVA Pairwise Tukey’s HSD post-hoc heatmaps for Accuracy, Kappa, Precision, and Recall, with positive mean differences (red) indicating CDI’s statistical advantage. **(d)** Model-wise precision boxplots reveal comparable precision across algorithms, confirming feature stability.

Supplementary Figure 9. Model-wise comparison of Featurizers by AUC-ROC, Accuracy, and Cohen’s Kappa. **(a)** Boxplots of AUC-ROC across nine classifiers (AdaBoost, CatBoost, Extra Trees, Gradient Boosting, LightGBM, Logistic Regression, Random Forest, SVM, XGBoost), showing CDI as the top or co-leading performer for nearly all models. **(b)** Accuracy boxplots showing CDI’s consistently higher mean values and reduced variance compared with deep and descriptor-based Featurizers. **(c)** Cohen’s Kappa boxplots illustrating strong inter-class agreement for CDI-based predictions, reaffirming feature consistency and balanced decision boundaries.

Supplementary Figure 10. Rank-based CDI performance across datasets and models. **(a)** Boxplots comparing F1-scores across nine classifiers, demonstrating CDI’s consistently higher medians and tight variance relative to competing Featurizers. **(b)** Boxplots of Balanced Accuracy values confirming CDI’s resilience against class imbalance and superior generalization across models. **(c)** Atom-count density plot of benchmark molecules (mean = 30.1, range = 1-384), confirming representative molecular size diversity in benchmarking datasets.

Supplementary Figure 11. CDI rank heatmaps across classification and regression models. **(a)** Heatmaps summarizing classification performance (AUC-ROC, Accuracy, Balanced Accuracy, F1 Score, Precision, Recall, Kappa) across datasets (TDC, BACE, PCBA, SIDER, Tox21, ToxCast). CDI consistently yields the top ranks across metrics and models, denoted by higher scoring. **(b)** Bar plots of Test Mean Absolute Error (MAE) for nine regression models (XGBoost, Random Forest, Gradient Boosting, Extra Trees, CatBoost, Lasso, and ElasticNet) showing CDI’s lowest average error, confirming cross-domain predictive strength.

Supplementary Figure 12. Comprehensive regression performance and rank-based analysis. **(a)** Heatmaps of regression metrics (R², MAE, RMSE, MSE) across ten physicochemical datasets showing CDI’s consistently high correlation and low error across models. **(b)** Median R² bar plots for nine regressors (Support Vector, Random Forest, Gradient Boosting, Extra Trees, XGBoost, CatBoost, Lasso, ElasticNet) demonstrating CDI’s dominant explanatory accuracy. **(c)** RMSE bar plots illustrating CDI’s minimal root-mean-square prediction error across all regression models, confirming superior stability and convergence. **(d)** MSE bar plots highlighting CDI’s lowest mean-squared error values relative to both deep learning and classical Featurizers, indicating high-accuracy regression performance.

Supplementary Figure 13. Visualization of molecular embeddings using t-SNE and PCA for chiral and kekulized datasets. **(a)** t-SNE projections for the chiral versus non-chiral dataset using embeddings generated by CDI, ChemBERTa, GROVER, ImageMol, MOPAC, Signaturizer, and Mordred. CDI and ChemBERTa exhibit the clearest visual separation between chiral (green) and non-chiral (pink) molecules, indicating enhanced stereochemical discrimination in their latent representations. **(b)** Quantitative clustering metrics (Silhouette Score, Davies-Bouldin Index, Calinski-Harabasz Score) for both t-SNE and PCA embeddings. CDI achieves the highest silhouette and Calinski–Harabasz values with the lowest Davies-Bouldin Index, confirming superior clustering quality and stereochemical sensitivity. **(c)** t-SNE projections for the kekulized versus non-kekulized SMILES dataset (orange vs. purple). CDI and ChemBERTa again demonstrate strong spatial segregation of the two classes, reflecting an improved capacity to encode electronic structure variations and aromatic bond alternation. **(d)** Quantitative comparison of clustering metrics for the kekulized dataset using t-SNE and PCA, confirming CDI’s top silhouette and Calinski-Harabasz scores among all Featurizers, signifying its highest discriminative fidelity for electronic structure encoding. **(e)** PCA parameter summary, listing principal component count (n_comp = 2), learning rate, iteration count (n_iter = 1000), and initialization method (“pca”) used for dimensionality reduction across all feature sets. **(f)** t-SNE parameter summary, detailing key hyperparameters including number of components (n_comp = 2), perplexity (42), random state, and iteration count (n_iter = 1000), ensuring reproducibility and comparability across embeddings. **(g)** Workflow schematic illustrating the cross-language validation procedure. A set of 100 representative SMILES molecules was processed through both the Python and R implementations of the Chemical Dice Integrator (CDI). Embeddings were generated independently from each environment, and pairwise cosine similarity was computed to assess embedding equivalence. **(h)** Cosine similarity plot showing per-SMILES comparison between R- and Python-derived CDI embeddings. The near-perfect similarity (mean cosine similarity > 0.99) confirms cross-platform reproducibility and ensures functional parity between CDI’s R and Python packages.

## SUPPLEMENTARY DATA

**Supplementary Table 1:** The table consists of packages and dependencies used in CDI development, including version numbers.

**Supplementary Table 2:** The table summarizes the 23 classification datasets with 171 tasks used for benchmarking CDI, including dataset names, sources, number of molecules, and endpoint types.

**Supplementary Table 3**: The table provides details of the 10 regression datasets employed in CDI benchmarking, specifying dataset sources and target properties.

**Supplementary Table 4**: The table presents comprehensive classification performance metrics for all Aggregator methods across datasets, reporting AUC-ROC, Balanced Accuracy, Precision, Recall, F1-score, and Cohen’s Kappa.

**Supplementary Table 5**: The table contains detailed regression performance metrics for all Aggregator methods across datasets, including R², MAE, RMSE, and MSE values.

**Supplementary Table 6**: The table shows statistical rankings of Aggregator methods for classification tasks, presenting mean ranks with 95% confidence intervals across datasets.

**Supplementary Table 7**: The table provides statistical rankings of Aggregator methods for regression tasks, including mean ranks and 95% confidence intervals, highlighting CDI’s top-tier performance.

**Supplementary Table 8**: The table includes comprehensive classification results for all molecular Featurizers-CDI, ChemBERTa, GROVER, ImageMol, Signaturizer, MOPAC, and Mordred across datasets, with performance metrics such as AUC-ROC, Balanced Accuracy, Precision, Recall, and F1-score.

**Supplementary Table 9**: The table summarizes regression results for all molecular Featurizers across 10 datasets, reporting R², MAE, RMSE, and MSE metrics to demonstrate comparative model performance.

**Supplementary Table 10**: The table reports the ranking of all molecular Featurizers on classification tasks with corresponding 95% confidence intervals, showing CDI’s consistent high-ranking performance.

**Supplementary Table 11**: The table lists the ranking of all molecular Featurizers for regression tasks with 95% confidence intervals, confirming CDI’s superior generalization and stability across datasets.

**Supplementary Table 12**: The table details computational resource benchmarking results, comparing runtime, CPU utilization, GPU memory usage, and total RAM consumption across all Featurizers under standardized Docker environments.

**Supplementary Table 13**: The table includes the dataset used for chiral versus non-chiral SMILES analysis, listing canonical SMILES, chiral center annotations, and stereochemical class labels used to test CDI’s sensitivity to stereochemical variation.

**Supplementary Table 14**: The table presents the dataset of canonical and kekulized SMILES used in electronic structure sensitivity analysis, providing both SMILES forms and molecular identifiers to assess CDI’s ability to distinguish bond-order variations

