## supplementary Figure 1 for "Chemical Dice Integrator (CDI): A Scalable Framework for Multimodal Molecular Representation Learning"

**a**

### Chemical Dice Integrator

1) Encoder Function generating latent represntations

$$\mathbf{h} = g(\boldsymbol{x})$$

2) Decoder reconstructspace from latent space

$$\tilde{\boldsymbol{x}} = f(\mathbf{h}) = f(g(\boldsymbol{x}))$$

3) Optimization objective minimizing reconstruction difference

$$\arg\min_{f,g} \quad \{(\Delta(\boldsymbol{x}), f(g(\boldsymbol{x}))\}$$

4) Dataset definition - each input has a embedding

$$D = \{\boldsymbol{x}^i \mid i = 1, \ldots, M\}, \quad \boldsymbol{x}^i = \{\, \boldsymbol{d}^i_j \mid j = 1, \ldots, 6\}$$

where M is total number of molecules

5) Mean Squared Error(MSE) loss function

$$L_{MSE}(x, y) = 1/n \sum_{i=1}^{n} |\boldsymbol{x}_i - \boldsymbol{y}_i|^2$$

where $\boldsymbol{x}, \boldsymbol{y} \in R^n$

6) Reconstruction Error (RE) to measure input-output differences

$$RE(x, \tilde{x}) = 1/n \sum_{i=1}^{n} |\boldsymbol{x} - \tilde{\boldsymbol{x}}_i|^2$$

where $\boldsymbol{x}, \tilde{\boldsymbol{x}} \in R^n$

7) ReLU activation function

$$ReLU(\boldsymbol{x}) = \max(\mathbf{0}, \boldsymbol{x})$$

8) Input and output dimension for each Semantic Commonality Autoencoder

$$\dim = \left[\sum_{k=1}^{6} |\mathbf{e}^i_k|\right] - |\mathbf{e}^i_j|$$

where|x| dimension of x

9) Concatenate all embeddings except jth one input to $j^{th}$ SCA block

$$\boldsymbol{a}^i_j = CONCAT(\boldsymbol{d}^i_{-j})$$

10) Latent embedding representation of $j^{th}$ for SCA encoder

$$\boldsymbol{p}^i_j = g(\boldsymbol{a}^i_j)$$

11) Combined SCA loss in MSE and RE terms

$$L_{SCAj} = L_{RE}(\boldsymbol{a}^i_j, \tilde{\boldsymbol{a}}^i_j) + L_{MSE}(\boldsymbol{p}^i_j, \boldsymbol{d}^i_j)$$

12) Super Embedding Autoencoder loss

$$L_{SEA} = L_{RE}(\boldsymbol{r}^i, \tilde{\boldsymbol{r}}^i)$$

13) Total Loss

$$L^i_{\text{total}} = \frac{1}{6} \sum_{j=1}^{6} L^i_{\text{SCA}_j} + L^i_{\text{SEA}}$$

**b**

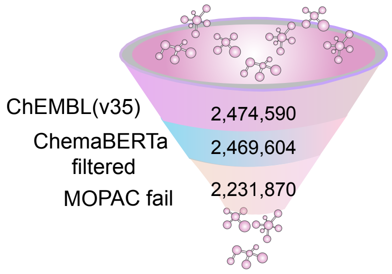


**c**

| Model | Basic | Generalised |
|---|---|---|
| Total Parameters | 1.09B | 0.10B |
| Learning Rate | 0.05 | 0.00001 |
| Training Size | 100% | 90% |
| Validation Size | 0% | 10% |
| Epoch | 600 | 30 |
| Total Molecule | 2231870 | 2231870 |
| Loss Criteria | MSE | MSE |
| Optimizer | SGD | AdamW |
| Scheduler | ReduceLROn Plateau | |
| Batch Size | 64 | 16 |

Supplementary Figure 1
