## Supplementary figures and images for "Chemical Dice Integrator (CDI): A Scalable Framework for Multimodal Molecular Representation Learning"

### supplementary Figure 2

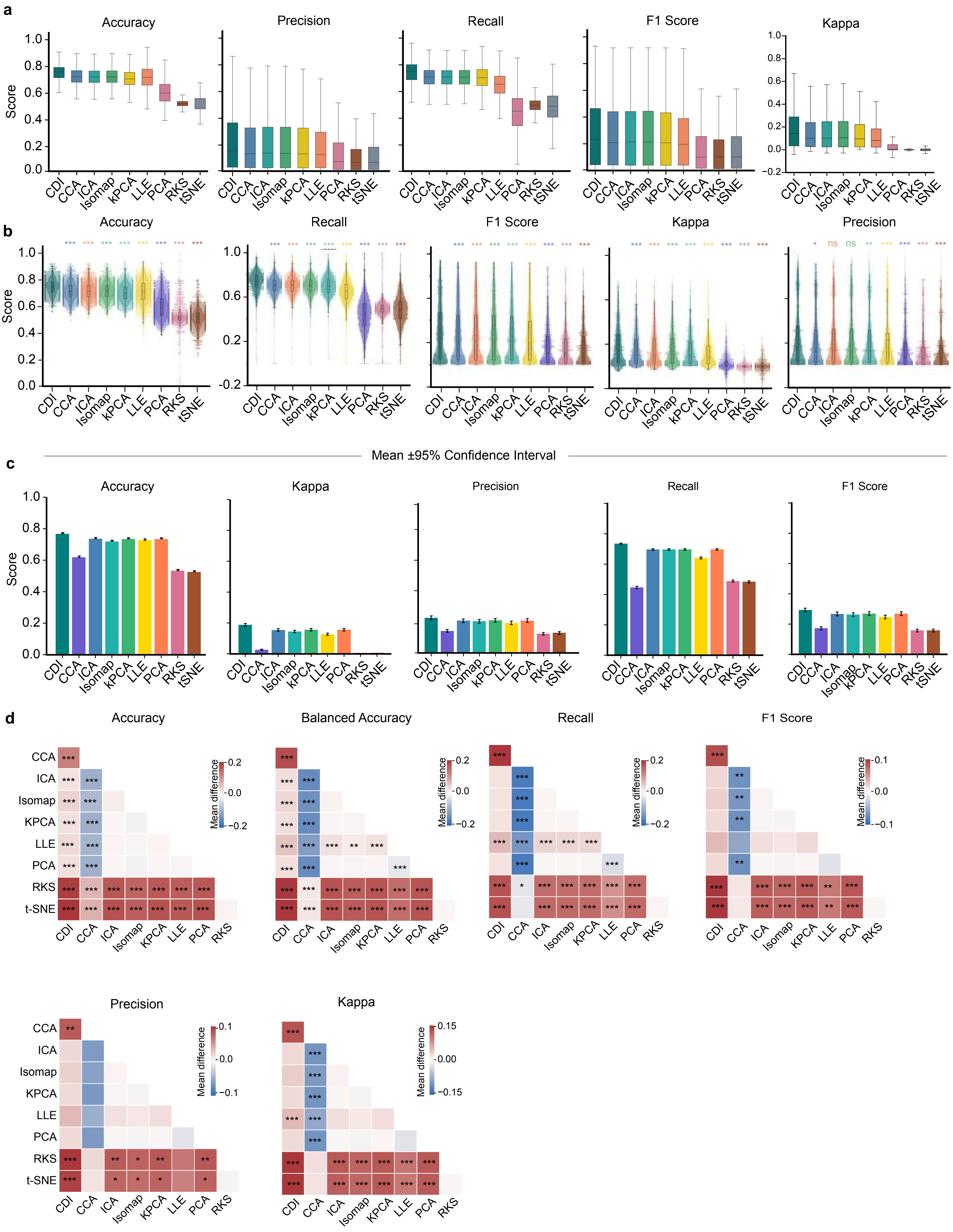


Supplementary Figure 2

### supplementary Figure 3

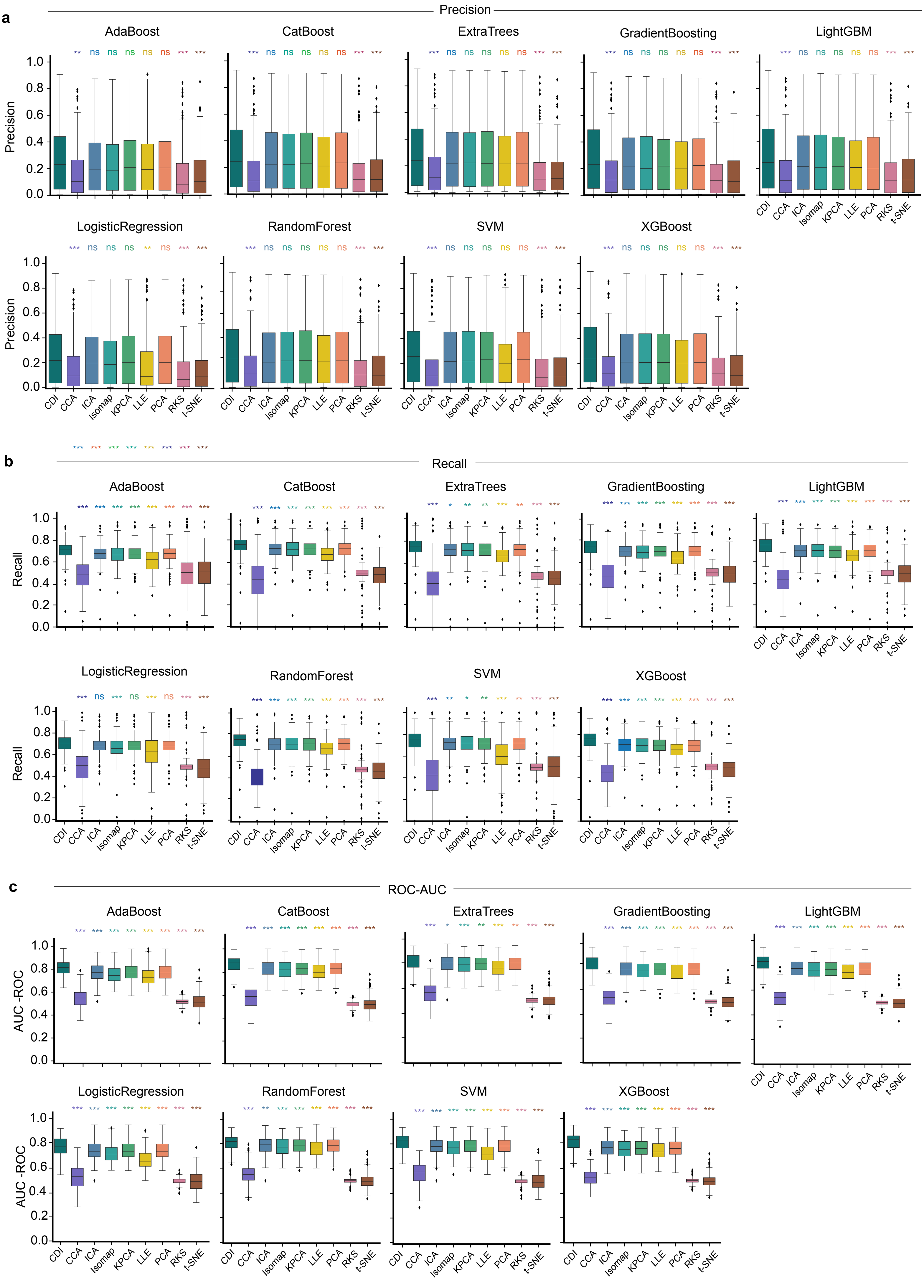


Supplementary Figure 3

### supplementary Figure 4

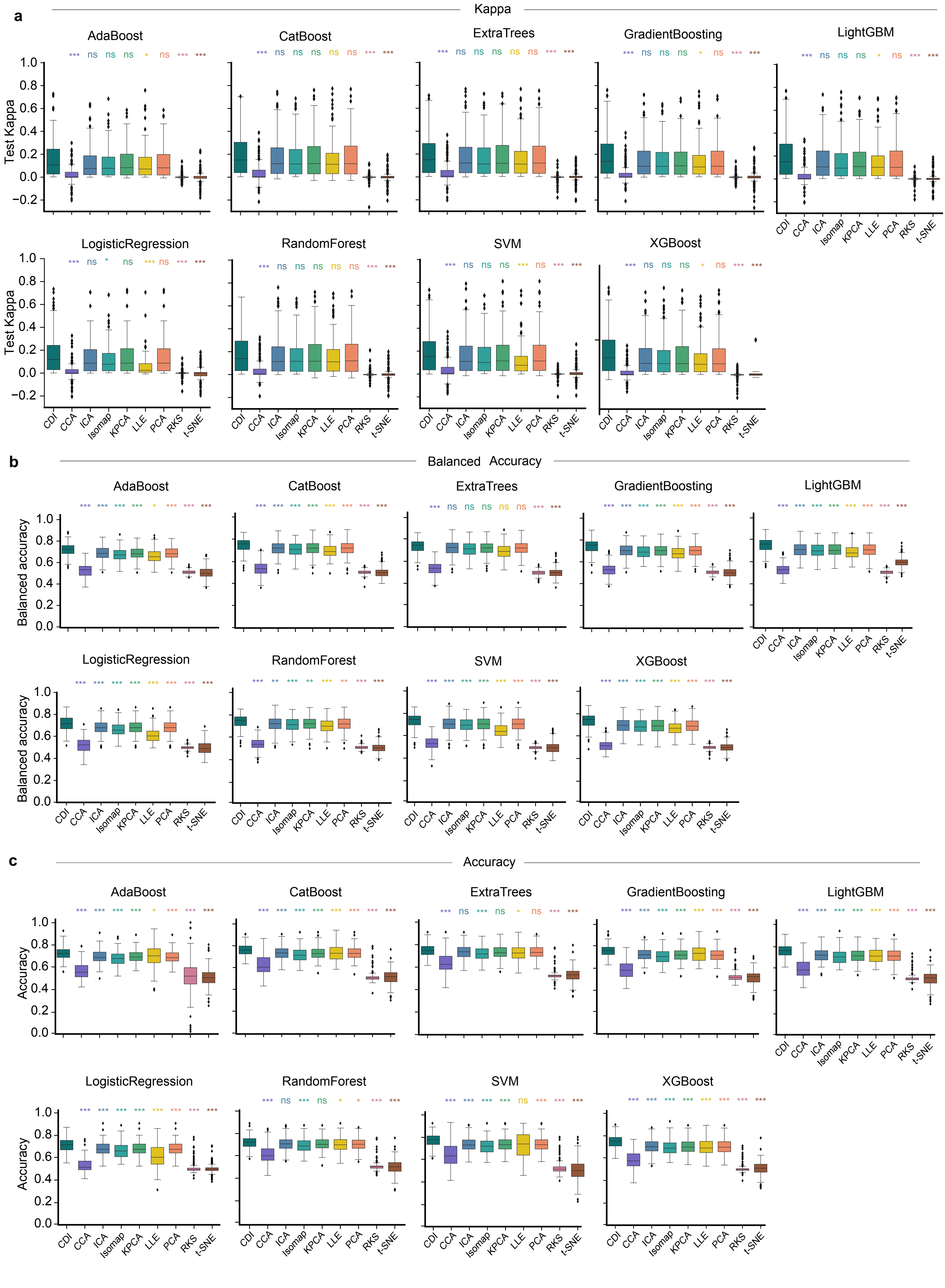


Supplementary Figure 4

### supplementary Figure 6

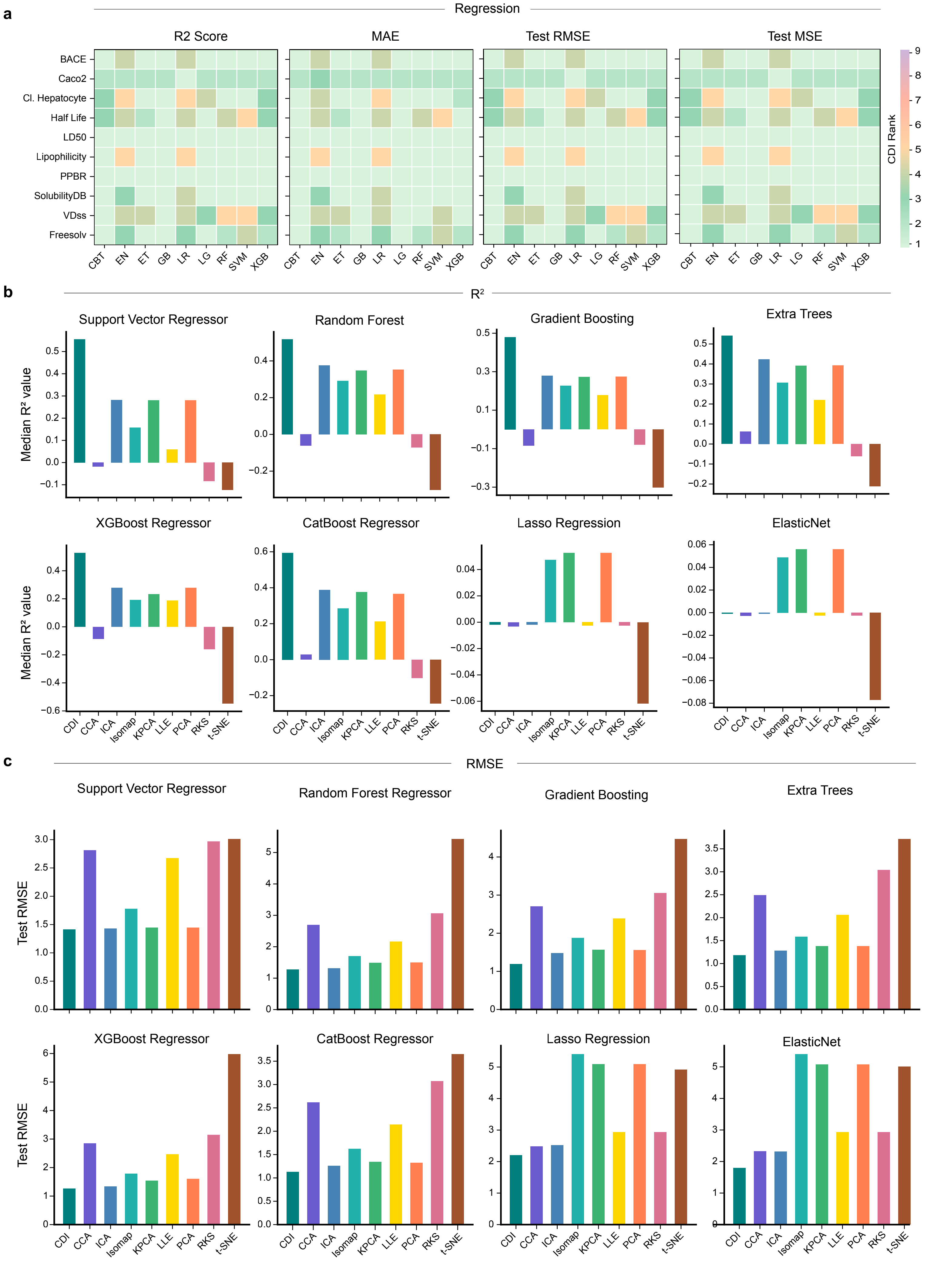


Supplementary Figure 6

### supplementary Figure 7

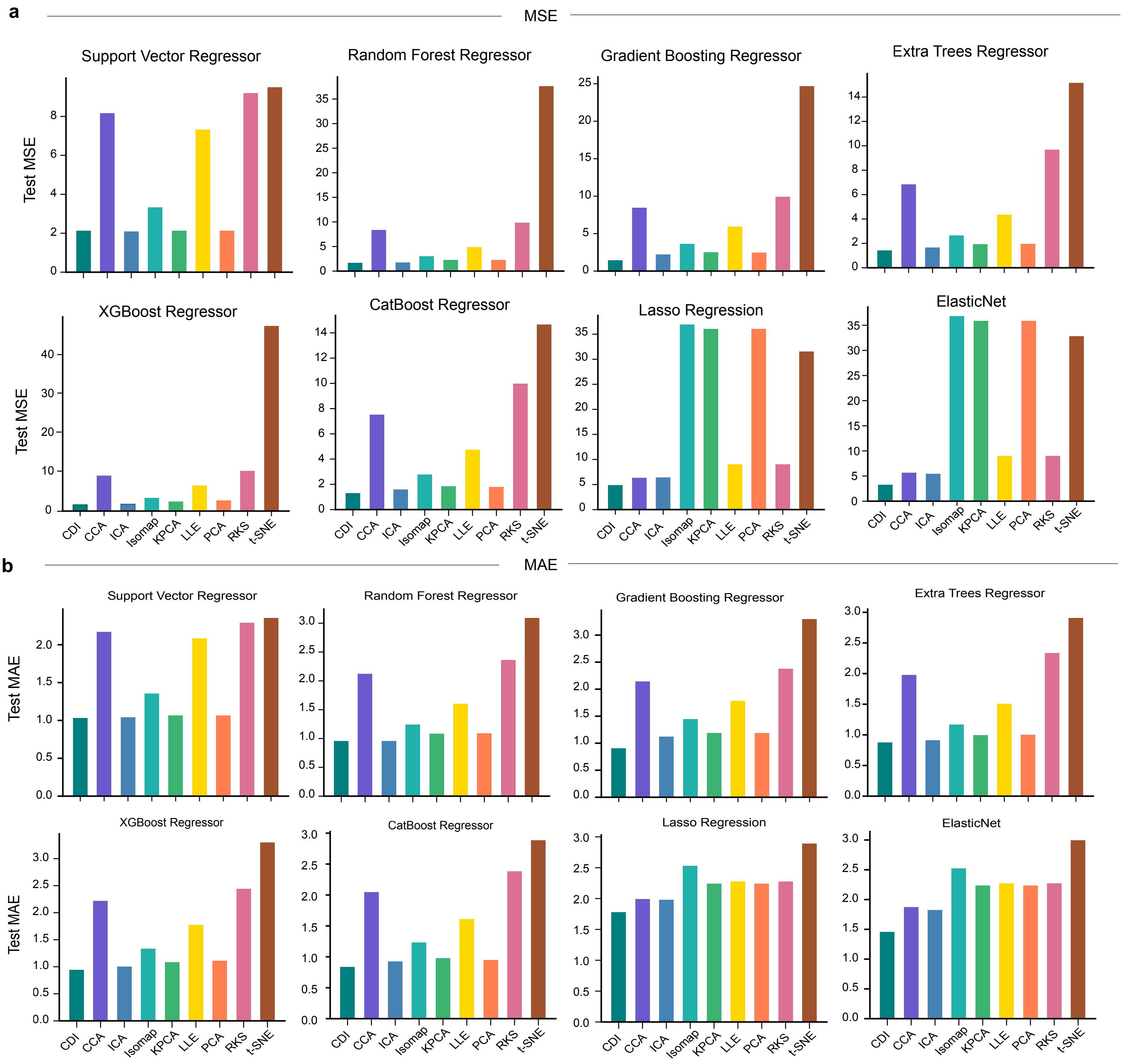


Supplementary Figure 7

### supplementary Figure 8

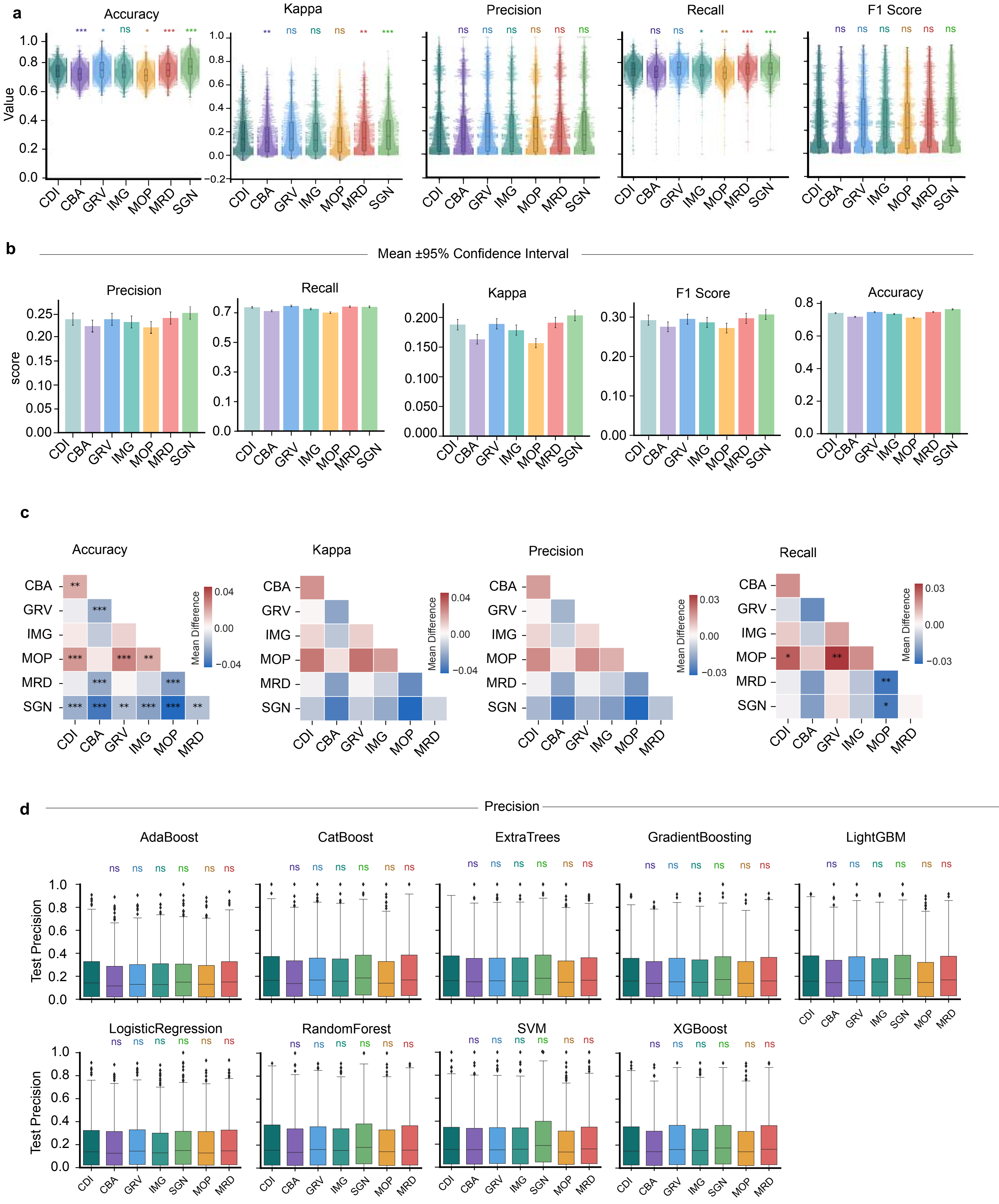


Supplementary Figure 8

### supplementary Figure 9

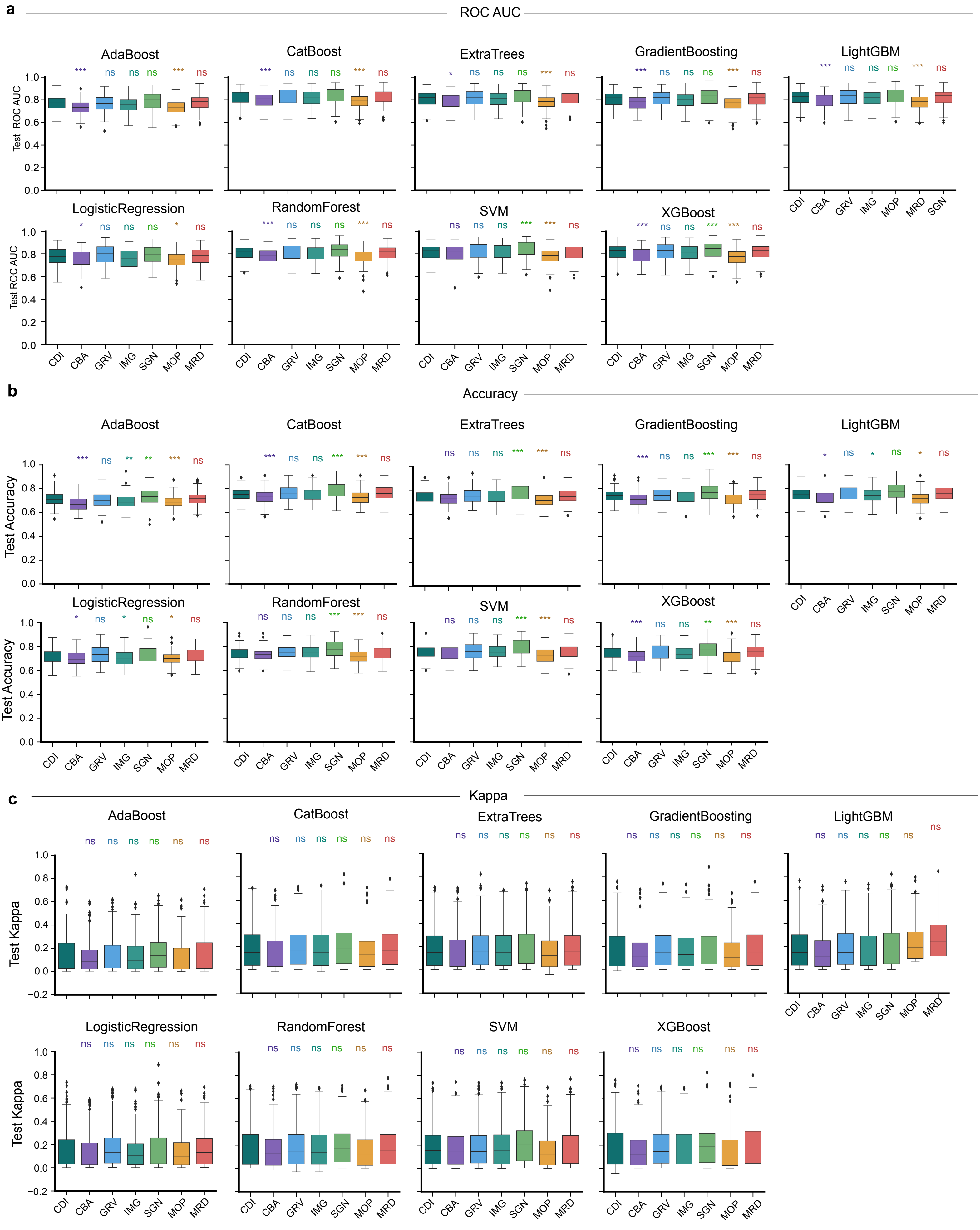


Supplementary Figure 9

### supplementary Figure 10

a

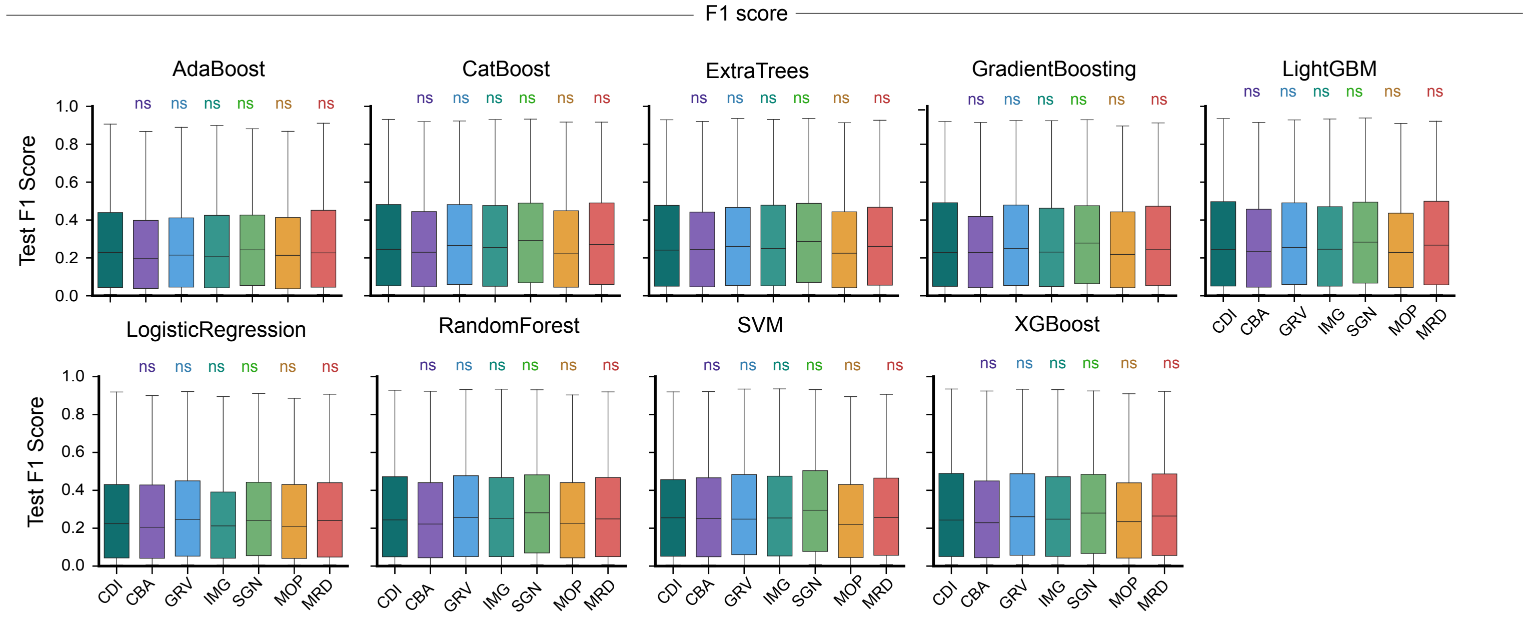


b

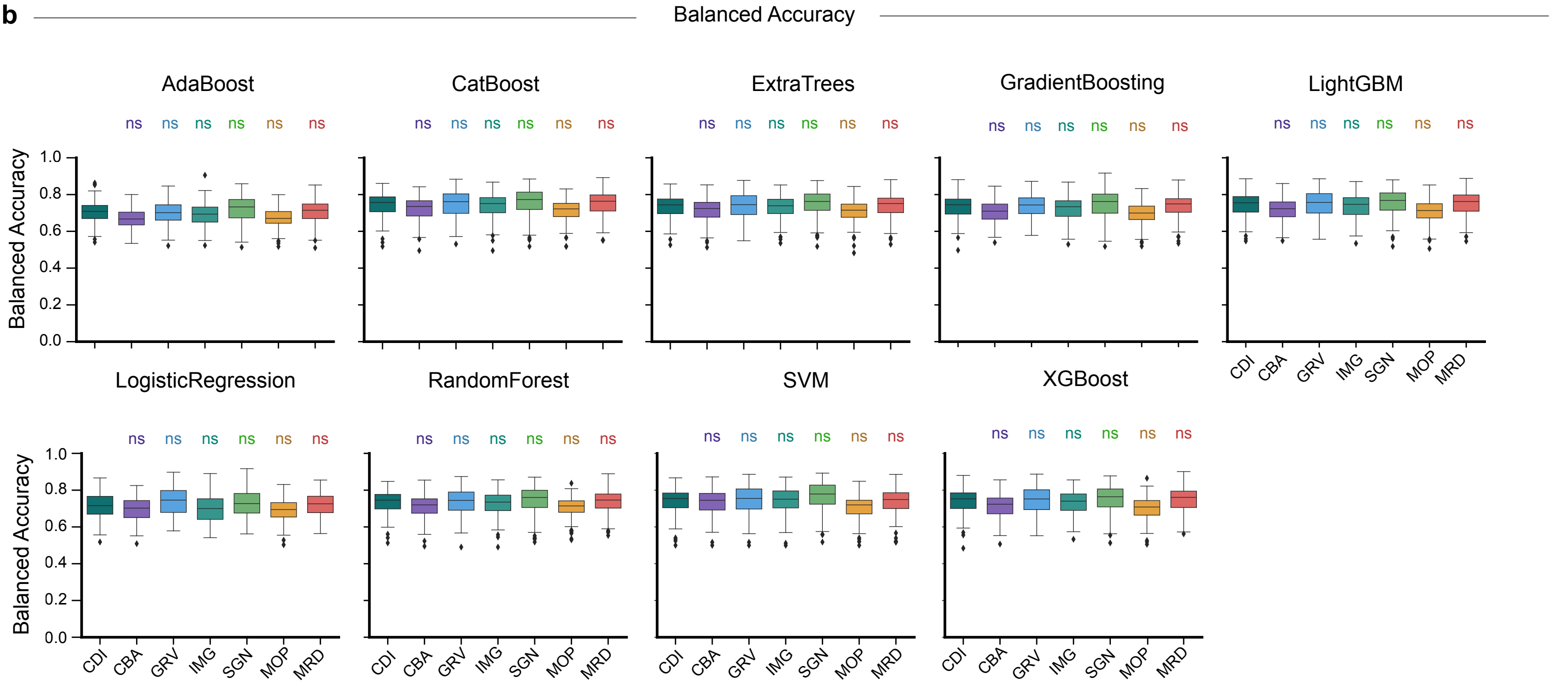


c

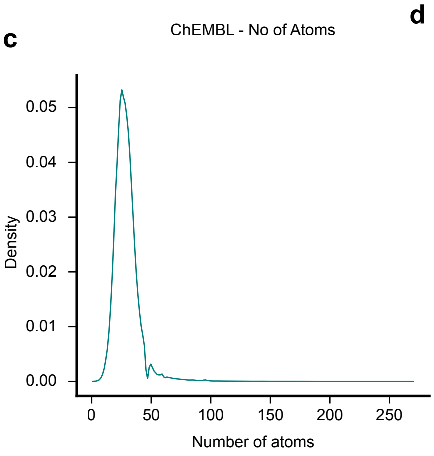


d

| Summary | Entity |
|---|---|
| Count | 2231870 |
| Mean | 29.046 |
| Std. | 10.682 |
| Min. | 1 |
| 25% | 23 |
| 50% | 28 |
| 75% | 33 |
| max | 287 |

Supplementary Figure 10

### supplementary Figure 11

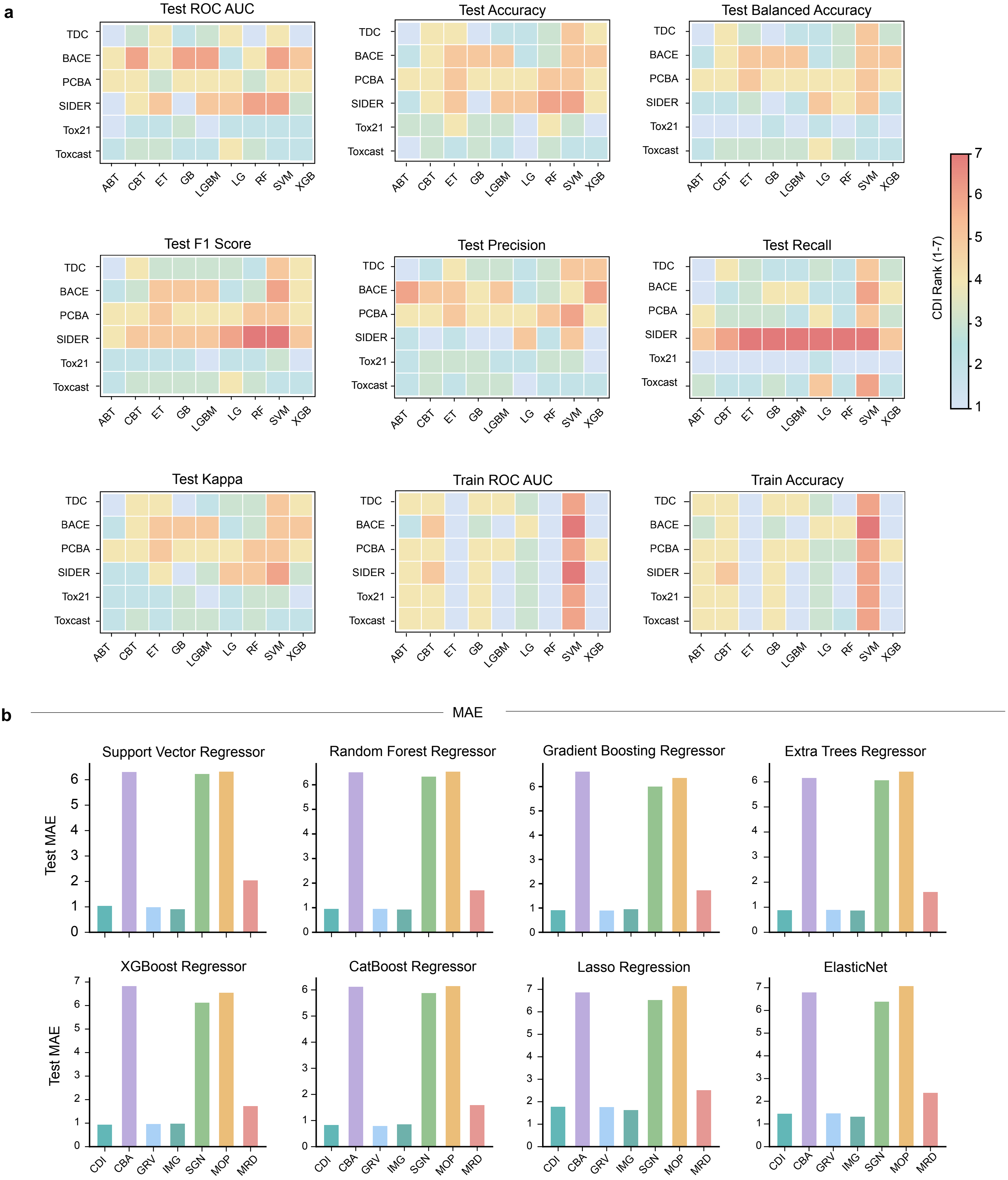


Supplementary Figure 11

### supplementary Figure 12

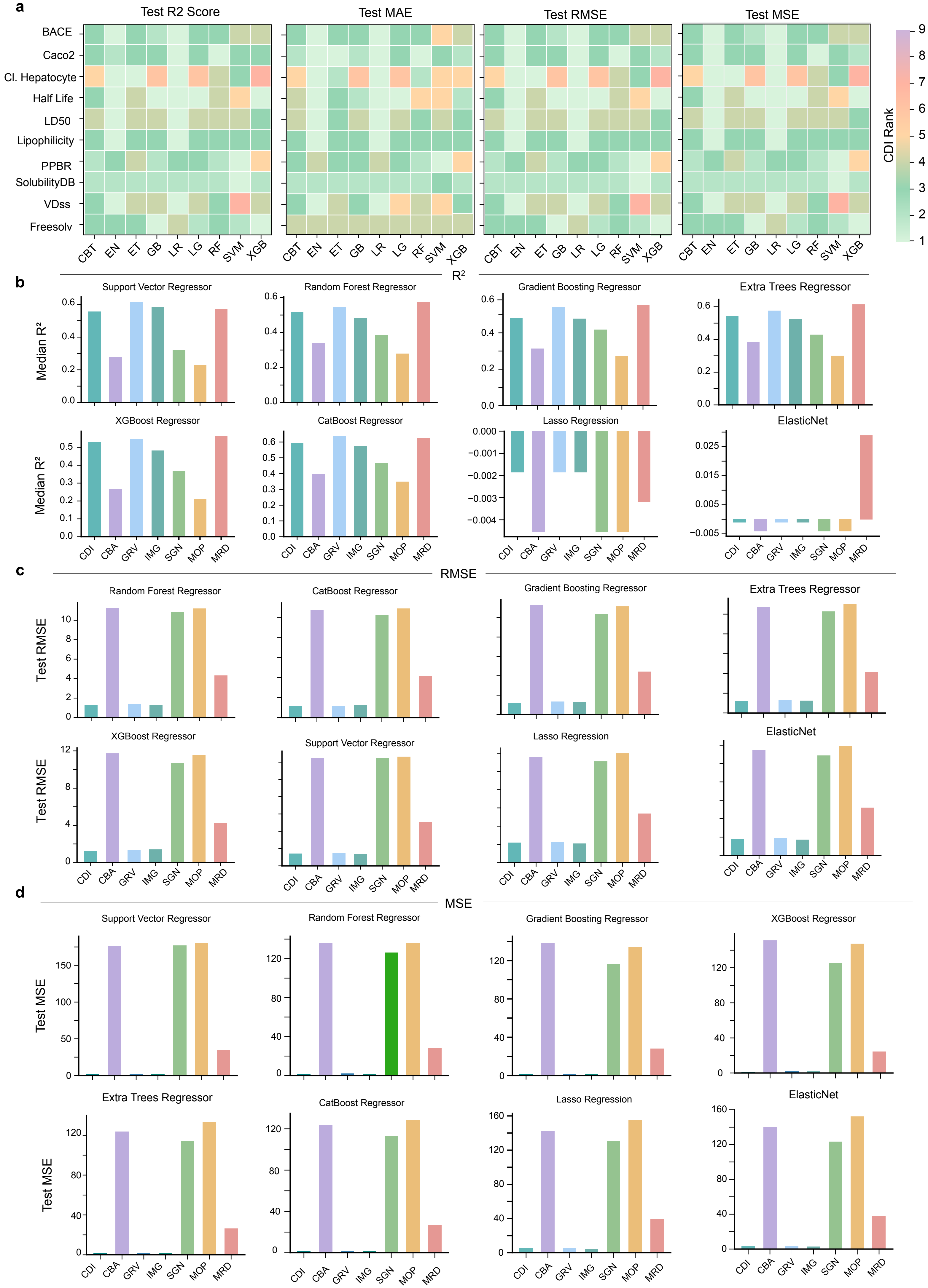


Supplementary Figure 12
