## supplementary Figure 5 for "Chemical Dice Integrator (CDI): A Scalable Framework for Multimodal Molecular Representation Learning"

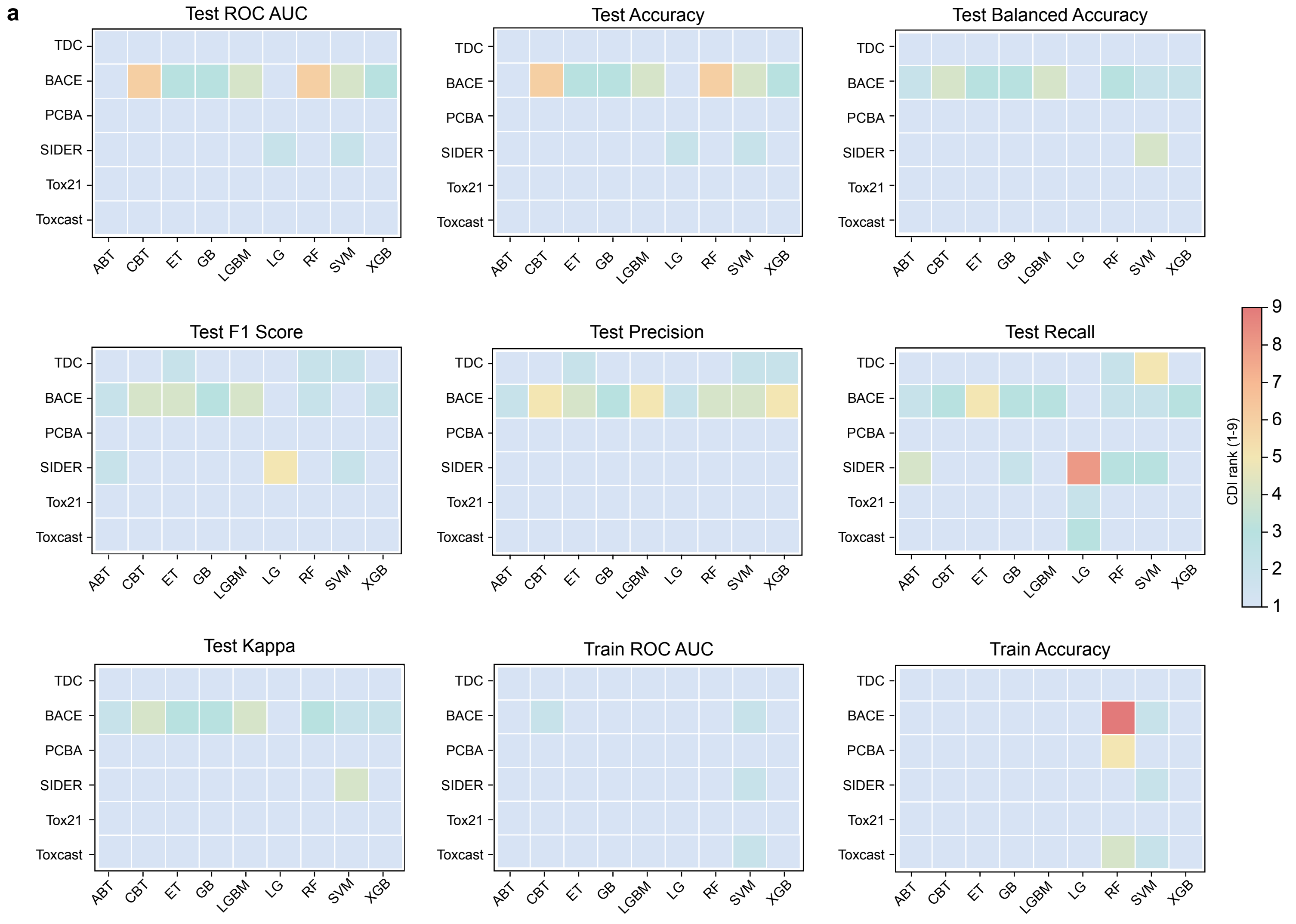


AdaBoost (ABT) CatBoost (CBT) ExtraTrees (ET) Gradient Boosting (GB) XGBoost (XGB) Light GBM (LGBM) Logistic Regression (LG) Random Forest (RF)

Supplementary Figure 5
