## supplementary Figure 13 for "Chemical Dice Integrator (CDI): A Scalable Framework for Multimodal Molecular Representation Learning"

**a**

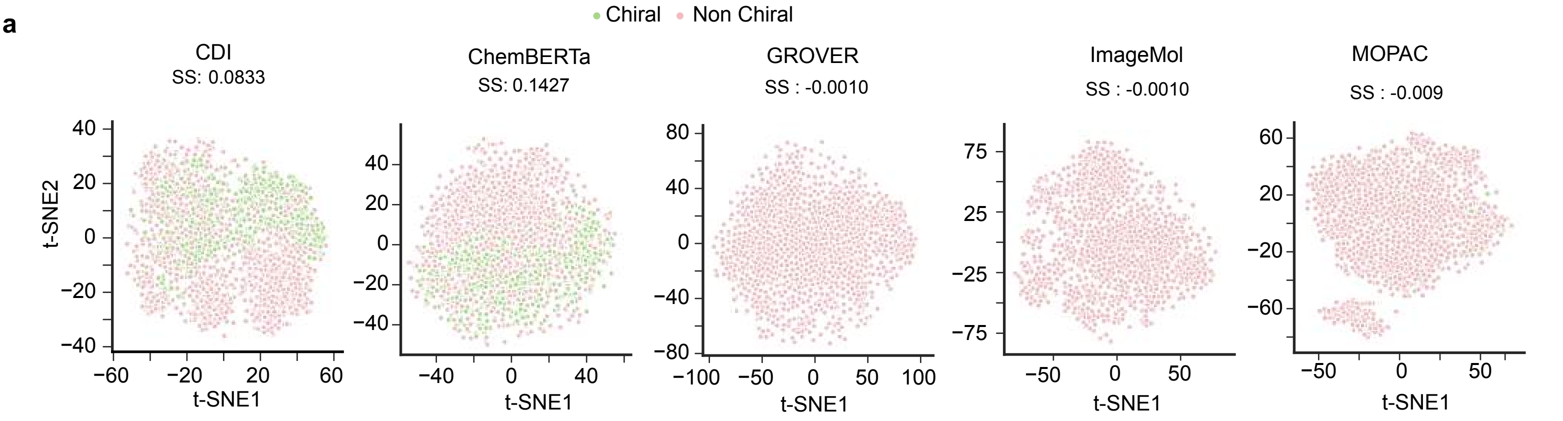


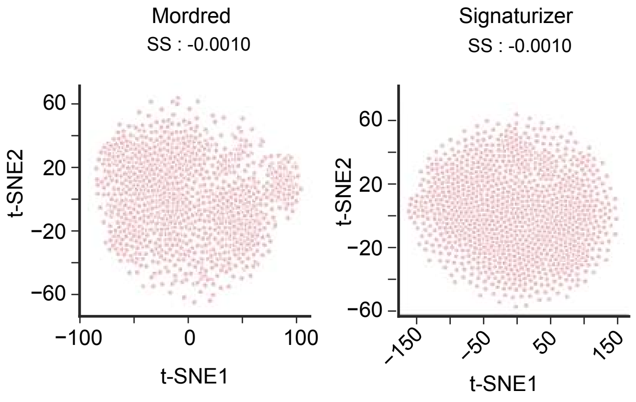


**b**

t-SNE

| Feature set | Silhouette | DB score | CH score |
|---|---|---|---|
| CDI | 0.0833 | 3.9984 | 105.016 |
| ChemBERTa | 0.1427 | 2.3704 | 294.6819 |
| GROVER | -0.0010 | 26852.28 | 0.0000 |
| ImageMol | -0.0010 | 1490.83 | 0.0008 |
| Signaturizer | -0.0010 | 0.0000 | 0.0000 |
| MOPAC | -0.009 | 32776.11 | 0.0487 |
| Mordred | -0.0010 | 186.7446 | 0.0000 |

PCA

| Feature set | Silhouette | DB score | CH score |
|---|---|---|---|
| CDI | 0.0518 | 4.2054 | 100.7358 |
| ChemBERTa | 0.0434 | 1.7710 | 92.1562 |
| GROVER | -0.0010 | 50789.68 | 0.0007 |
| ImageMol | -0.0009 | 345.5906 | 0.0555 |
| Signaturizer | -0.0010 | 0.0000 | 2.1843 |
| MOPAC | -0.0008 | 82.916 | 2.102 |
| Mordred | -0.0009 | 152.033 | 0.0695 |

**c**

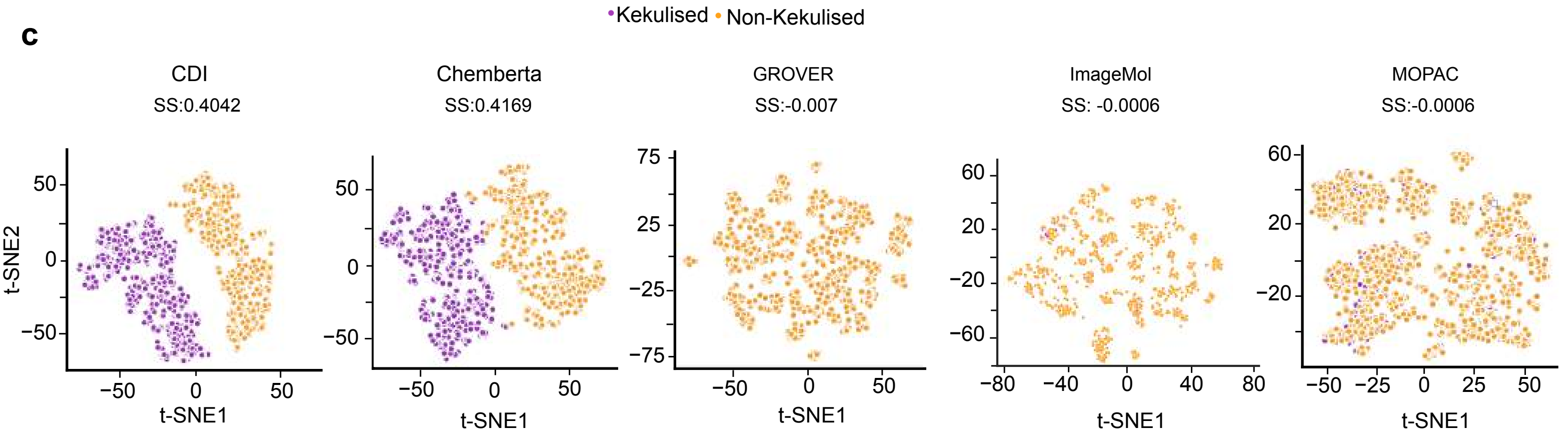


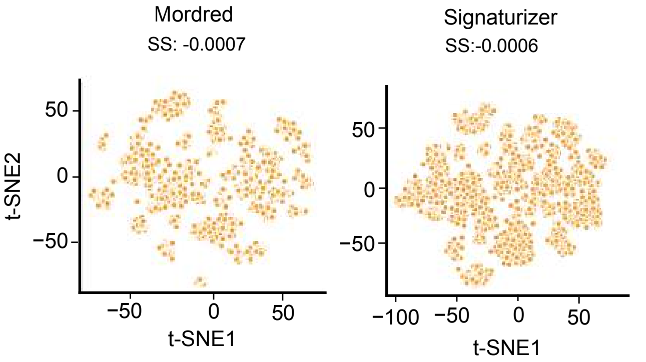


**d**

t-SNE

| Feature set | Silhouette | DB score | CH score |
|---|---|---|---|
| CDI | 0.4042 | 1.1128 | 2073.073 |
| ChemBERTa | 0.4169 | 1.0257 | 2510.446 |
| GROVER | -0.0007 | 577548 | 0.0000 |
| ImageMol | -0.0006 | 3049.29 | 0.0003 |
| Signaturizer | -0.0007 | 26890.08 | 0.0000 |
| MOPAC | -0.0006 | 560.12 | 0.0087 |
| Mordred | -0.0007 | 4771.94 | 0.0001 |

PCA

| Feature set | Silhouette | DB score | CH score |
|---|---|---|---|
| CDI | 0.6066 | 0.6539 | 4364.88 |
| ChemBERTa | 0.5681 | 0.7069 | 4786.15 |
| GROVER | -0.0007 | 7178.095 | 0.0000 |
| ImageMol | -0.0006 | 236.59 | 0.0495 |
| Signaturizer | -0.0007 | | 0.000 |
| MOPAC | -0.0005 | 277.967 | 0.0329 |
| Mordred | -0.0006 | 125.233 | 0.1557 |

**e**

PCA

| Parameter | Value |
|---|---|
| n_comp | 2 |
| svd_solver | auto |
| random_state | 42 |
| iterated_power | auto |

**f**

t-SNE

| Parameter | Value |
|---|---|
| n_comp | 2 |
| random_state | 42 |
| perplexity | 30 |
| n_iter | 1000 |
| init | pca |
| learning rate | auto |

**g**

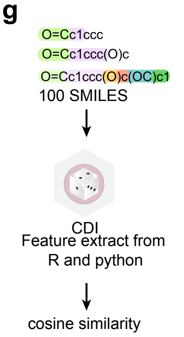


**h**

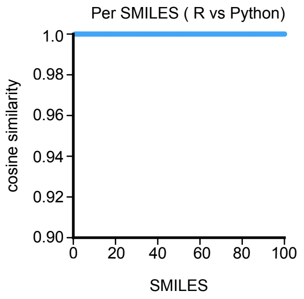


Supplementary Figure 13
